# Anvaya: phylogenomic analysis of chromosome fusion-and-mixing characters and detection of shared whole-genome duplication

**DOI:** 10.64898/2026.10.08.757583

**Authors:** Siddharth Kulkarni

**Affiliations:** CSIR-Center for Cellular and Molecular Biology, Hyderabad, India; Academy of Scientific and Innovative Research (AcSIR), Ghaziabad, India

**Keywords:** macrosynteny, ancestral linkage groups, chromosome fusion, whole-genome duplication, genome architecture evolution, phylogenetic software

## Abstract

Chromosome-scale conservation of gene linkage (‘macrosynteny’) provides phylogenetic characters largely independent of sequence data. Fusion of ancestral linkage groups, and especially the subsequent interspersion (“mixing”) of their genes, is argued to be effectively irreversible, and such synapomorphic events have been used to resolve deep, contentious nodes. I present Anvaya, a dependency-light, open-source toolkit of two Python programs with companion threshold-calibration tools which takes reciprocal-best-hit (RBH) synteny data directly to a phylogenetic analysis that codes fusion and mixing as discrete characters under explicit optimality criteria. The synteny_builder module builds RBH orthology tables from genome assemblies, using either a designated reference genome or a multi-genome consensus procedure that reconstructs linkage groups de novo, requiring no curated reference set. The character_maps module codes every pairwise linkage-group co-occurrence as unfused, fused-clean or fused-and-mixed from gene order and a Fisher’s exact test, then searches for the most parsimonious trees under an unordered (Fitch/Sankoff) and an ordered, irreversible (Camin–Sokal) cost model, optionally under a backbone constraint. Every tree is accompanied by ancestral states, synapomorphies mapped onto clades, consistency and retention indices, bootstrap support, and a permutation test of phylogenetic signal. Anvaya also detects whole-genome and segmental duplication from an intragenomic comparison and recovers the ohnologs that reciprocal best hit structurally drops, which can otherwise render a retained linkage group invisible. Outputs include vector figures and matrices in NEXUS and TNT/Hennig86 formats for verification in PAUP* and TNT. On synthetic data, consensus reconstruction resists a conflicting genome, rescue reverses a total false negative, and the diagnostics flag a deliberately homoplastic character. Across thirteen acarine and outgroup genomes, weakly supported associations concentrate in the most distant genome and in poorly assembled regions. In two duplicated chelicerates, ohnolog rescue recovered 12.5% (*Limulus polyphemus*) and 2.8% (*Mastigoproctus giganteus*) and restored three *Limulus* chromosomes that otherwise carried none.

## INTRODUCTION

Resolving the deepest nodes of the tree of life has been a story of successive data types, each extending resolution beyond the last. Transcriptomic and other phylogenomic matrices displaced single-locus studies and settled many inter-phyletic and ordinal relationships, but left a residue of recalcitrant nodes that additional sequence data did not resolve (Jeffroy et al., 2006; Philippe et al., 2011; Kulkarni et al., 2021). Targeted enrichment of ultraconserved elements (UCEs) then made dense taxon sampling affordable, including from museum material. UCEs are loci conserved across distantly related taxa, and their variable flanking sequence is informative at shallower levels (Faircloth et al., 2012). Lineage-specific probe sets proved decisive where generic baits captured too few loci, in diversity rich groups ranging from Hymenoptera (Faircloth et al., 2015) to spiders (Kulkarni et al., 2020), Opiliones (Derkarabetian et al. 2023) and Chelicerata more broadly (Bhoi et al., 2026). Most recently, chromosome-level assemblies have opened another phase, in which genome architecture itself, rather than aligned sequence, becomes the comparative unit (Schultz et al., 2023; Kulkarni et al., 2024; Klementz et al., 2025; Vargas-Chávez et al., 2025).

Certain nodes nonetheless remain obdurate, and the reasons are increasingly well characterised. Rapid radiations compress the internodes across which substitutions accumulate. Even very large matrices therefore carry few informative changes per node, while retaining ample opportunity for conflicting signals (Rokas & Carroll, 2006; Whitfield & Lockhart, 2007). Elevated substitution rates and regressive morphological evolution, both common in miniaturised lineages such as Acariformes (mites), compound the problem by generating homoplasy that scales with the data rather than washing out. Bursts of diversification leave short, deep internodes flanked by long terminal branches, the configuration in which systematic error is most readily mistaken for signal (Felsenstein, 1978; Philippe et al., 2011). Adding sequence data under these conditions has been observed to increase nodal support without increasing accuracy. The recalcitrant interrelationships within the class Arachnida are a case in point. Despite successive phylogenomic datasets, many ordinal relationships remain contested, as does the monophyly of Acari (Sharma et al., 2014; Ballesteros et al., 2022; Domènech et al. 2026).

This is the setting in which rare genomic changes are valuable. Characters such as gene-cluster rearrangement, intron gain and loss, and chromosome-scale fusion arise infrequently and are unlikely to be acquired independently. They are also largely decoupled from the substitutional processes that generate long-branch artefacts, so they provide an additional line of robust evidence that is not simply more of the same (Rokas & Holland, 2000; Boore, 2006; Klementz et al., 2025). Macrosynteny is among the most tractable of these characters. Fusion-with-mixing of genes in particular has been argued to be effectively irreversible once established, which gives it known polarity (Schultz et al., 2023), however, limits of this data class and interpretation are being gradually realized. Rearrangement is far more pervasive across bilaterians than once assumed (Lewin et al., 2025b). Macrosyntenic characters can recapitulate rather than resolve existing conflict (Plessy et al., 2024; Kulkarni et al., 2026), and the statistical basis for calling them has been questioned (Copley, 2025). Both considerations warrant support for a pipeline in which every threshold is explicit and adjustable, and in which homoplasy is measured rather than assumed away.

Whole-genome duplication is a rare genomic change of a particular kind where a duplicated genome is inherited by every descendant lineage. Therefore, the ancestral duplication and its architecture shared by a group of taxa are itself a synapomorphy, and unlikely to arise again in the same form (Ohno, 1970). Ancient duplications have been placed on the vertebrate stem (Dehal & Boore, 2005; Simakov et al., 2020) and at the base of teleost fishes (Jaillon et al., 2004). Among chelicerates, one occurred in the ancestor of Arachnopulmonata, the clade uniting spiders, scorpions and their relatives (Schwager et al., 2017), and duplication has occurred separately thrice in horseshoe crabs (Kenny et al., 2016; Nong et al., 2021). The arachnopulmonate duplication has been used explicitly as phylogenetic evidence which helped place pseudoscorpions within Arachnopulmonata where sequence-based analyses were compromised by long-branch attraction (Ontano et al., 2021). Its signature is macrosyntenic: a shared duplication leaves each ancestral linkage group represented on two ohnologous chromosomes, a pattern readable directly from chromosome-level assemblies (Kulkarni et al., 2024). Yet this is precisely the pattern that standard synteny pipelines are poorly equipped to detect, as I describe below.

In recent years, comparative genomics has converged on a striking regularity: many animal genomes retain chromosome-scale blocks of orthologous genes that have stayed physically linked since before the divergence of major animal lineages. This principle of conserved ancestral linkage groups (ALGs) was established for chordates. Seventeen linkage groups are conserved between amphioxus and vertebrates, which showed that syntenically equivalent chromosomes in distantly related lineages must reflect chromosomes present in their common ancestor (Putnam et al., 2008). The idea was formalised as a probabilistic model and used to reconstruct ancestral vertebrate genomes (Nakatani & McLysaght, 2017; Nakatani et al., 2021). It was then extended to sponges and cnidarians, defining ancestral bilaterian and metazoan linkage groups traceable across the animal tree (Simakov et al., 2020, 2022). However, some lineages have eroded this ancestral synteny and rewired their genomes to establish linkage groups of their own. Nematodes and drosophilids are examples, with their Nigon and Muller elements respectively (Tandonnet et al., 2019; Simakov et al., 2022; Ivanković et al., 2024), Lepidoptera, with their ancestral Merian elements (Wright et al., 2024). Across bilaterians more broadly, conservation proves to be the exception rather than the rule. Most sampled genomes, including every ecdysozoan examined, are highly rearranged (Lewin et al., 2025b). Massive lineage-specific restructuring is documented in annelids (Lewin et al., 2024; Vargas-Chávez et al., 2025), bryozoans (Lewin et al., 2025a) and nematodes (Simmons et al., 2024).

Because these patterns are set by rare chromosomal fusion, fission and translocation rather than by nucleotide substitution, they provide characters largely orthogonal to sequence alignment. Fusion, once it has occurred, is rarely undone. Across Lepidoptera fusions are rare and fissions are rarer, and chromosomes derived from ancient fusions retain syntenic domains that reflect the original elements (Wright et al., 2024). Where two fused linkage groups have become interleaved, that mixing is a one-way process: its reversal is about as probable as two mixed fluids spontaneously separating (Schultz et al., 2023). Fusion-with-mixing, coded as a discrete, polarity-known, non-reversible character, has been used to resolve one of the most contentious nodes in the animal tree. Fusion-with-mixing events shared by sponge, cnidarian and bilaterian genomes placed ctenophores, rather than sponges, as sister to all other animals (Schultz et al., 2023). Thereby, one of the most consequential recent results in deep animal phylogenetics was thus, at bottom, a cladistic argument: discrete characters, explicit polarity, and a parsimony criterion matched to how those characters are believed to evolve.

The computational infrastructure for this style of analysis is only gradually catching up with the idea. The most widely used tool, odp, computes reciprocal-best-hit (RBH) orthology between a query genome and a reference ALG set. It tests each chromosome–ALG pairing for enrichment with Fisher’s exact test, and draws the Oxford dot plots and ribbon diagrams that have become standard for these data (Schultz et al., 2023). Several other tools have followed, briefly: (1) macrosyntR tests pairwise chromosome associations and clusters the chromosomes derived from the same ALG (El Hilali & Copley, 2023); (2) Syngraph reconstructs ancestral linkage groups node by node across a phylogeny, using a fusion– fission model (Mackintosh et al., 2023); (3) LINGUINE reconstructs them using orthogroups, a hidden Markov model and explicit collapse of duplicated linkage groups (Vargas-Chávez & Fernández, 2026); (4) egt quantifies continuous measures of genome mixing and estimates rates of change along an already specified, calibrated tree (Schultz & Simakov, 2026). Each of these tools takes the phylogeny as given, or compares genomes pairwise; none infers it. What is missing is the complementary step: a program that codes fusion and mixing as discrete, ordered character states and runs a parsimony tree search on them, as a cladist would analyse a morphological matrix. It should apply alternative, clearly stated cost models and allow a fixed backbone hypothesis to be tested. Its output should also be verifiable independently in TNT or PAUP* (Swofford, 2003; Goloboff et al., 2008). Moreover, the curated ALG reference sets now available, such as the Bilaterian–Cnidarian–Sponge (BCnS) ALGs (Simakov et al., 2022), were built for deep phylogenetic questions. Given how extensively genomes are rearranged in many clades, they are not necessarily an appropriate yardstick for shallower questions, within an order or family, where no equivalent curated reference exists.

There is also an open question of how the statistical significance of an ALG–chromosome association should be assessed. Copley (2025) argued that the permutation procedure used to assign significance to candidate ALGs treats all taxa as equally distant, and so can overestimate significance, while some real linkage groups may fail to reach significance under the same procedure. While the critique does not overturn the biological argument for fusion-with-mixing as an informative, irreversible character, it does warrant a software in which the significance criterion, and every other threshold used to call a character state, is an explicit, adjustable parameter rather than being a fixed constant.

Whole-genome duplication raises a further, largely unaddressed problem. Reciprocal best hit is a strict one-to-one mapping by construction (Tatusov et al., 1997; Moreno-Hagelsieb & Latimer, 2008). Suppose a genome carrying a lineage-specific duplication, such as a spider descended from the arachnopulmonate duplication, is compared with a non-duplicated outgroup such as a tick. One member of every pair of duplicate-derived orthologs (ohnologs) is then silently dropped from the orthology table (Catchen et al., 2009). The result is not the doubled hit pattern one might hope to use as a diagnostic, but its opposite. Orthologs are divided between the two retained copies of each ancestral chromosome, neither copy may clear the threshold for calling an association, and a fully retained linkage group can disappear from the analysis altogether. A shared duplication, which should be among the most conspicuous characters available, is thereby rendered invisible. Collapsing duplicated linkage groups before ancestral reconstruction avoids artefacts where duplication is a confounder (Vargas-Chávez & Fernández, 2026). However, it discards the very signal of interest when the duplication itself is the question.

Here, I describe Anvaya, an open-source toolkit of two command-line Python programs with companion tools for threshold calibration, which contributes:

1. coding of pairwise linkage-group co-occurrence as a three-state ordered character (unfused, fused-clean, fused-and-mixed) directly from RBH ortholog tables, with every threshold adjustable;
2. a Sankoff-based parsimony search under unordered (Fitch-equivalent) and ordered, strictly irreversible (Camin–Sokal) cost models (Camin & Sokal, 1965; Fitch, 1971; Sankoff, 1975), exhaustive for small taxon samples and NNI-heuristic otherwise, with optional backbone constraints;
3. ancestral-state reconstruction and automatic mapping of the synapomorphies supporting each clade;
4. consensus reconstruction of linkage groups de novo from user-chosen core genomes, removing dependence on a single reference or a curated ALG set, with a faster single-reference mode and a sweep tool for choosing the thresholds;
5. detection of lineage-specific or shared whole-genome duplication by intragenomic self-comparison, and recovery of the ohnologs reciprocal best hit drops, with per-taxon thresholds calibrated against karyotype expectation rather than significance alone;
6. diagnostics reported with every tree: consistency and retention indices (Kluge & Farris, 1969; Farris, 1989), bootstrap support, a permutation test of phylogenetic signal, branch rate models, and a report on character non-independence from compound fusions;
7. assembly-contiguity reporting for every input genome, and export to NEXUS and TNT/Hennig86 for independent verification; and
8. publication-ready vector figures with editable text: Oxford dot plots, ideograms, character-matrix heatmaps, cladograms with synapomorphies on branches, ribbon plots with tree-derived track order and crossing-minimised chromosome order, and duplication views.

The Methods describe the design and algorithms; the toolkit is validated on synthetic data and applied to thirteen acarine and outgroup genomes and to two duplicated chelicerate genomes, with an unduplicated tick for comparison.

### Software description

#### Overview and availability

Anvaya comprises two command-line programs, genomes_to_rbh.py and alg_parsimony.py, and three companion tools for calibrating thresholds. All are implemented in Python 3 with pandas as the only hard dependency; matplotlib, SciPy and Biopython are used for figures, enrichment tests and coding-sequence translation respectively, and the software degrades with an explicit warning, when they are absent. genomes_to_rbh.py takes genome assemblies and writes reciprocal-best-hit (.rbh) ortholog tables. alg_parsimony.py takes .rbh tables, whether produced by genomes_to_rbh.py, by odp (Schultz et al., 2023) or by any pipeline emitting the same column layout, and produces the character matrix, trees, ancestral states, synapomorphy reports, diagnostics and figures. The companion tools choose thresholds from the data: sweep_consensus_params.py for linkage-group reconstruction, calibrate_wgd.py for duplication detection, and make_toy_dataset.py, which generates a simulated synthetic dataset with known ground truth for testing (Fig. 1).

**Figure 1.**
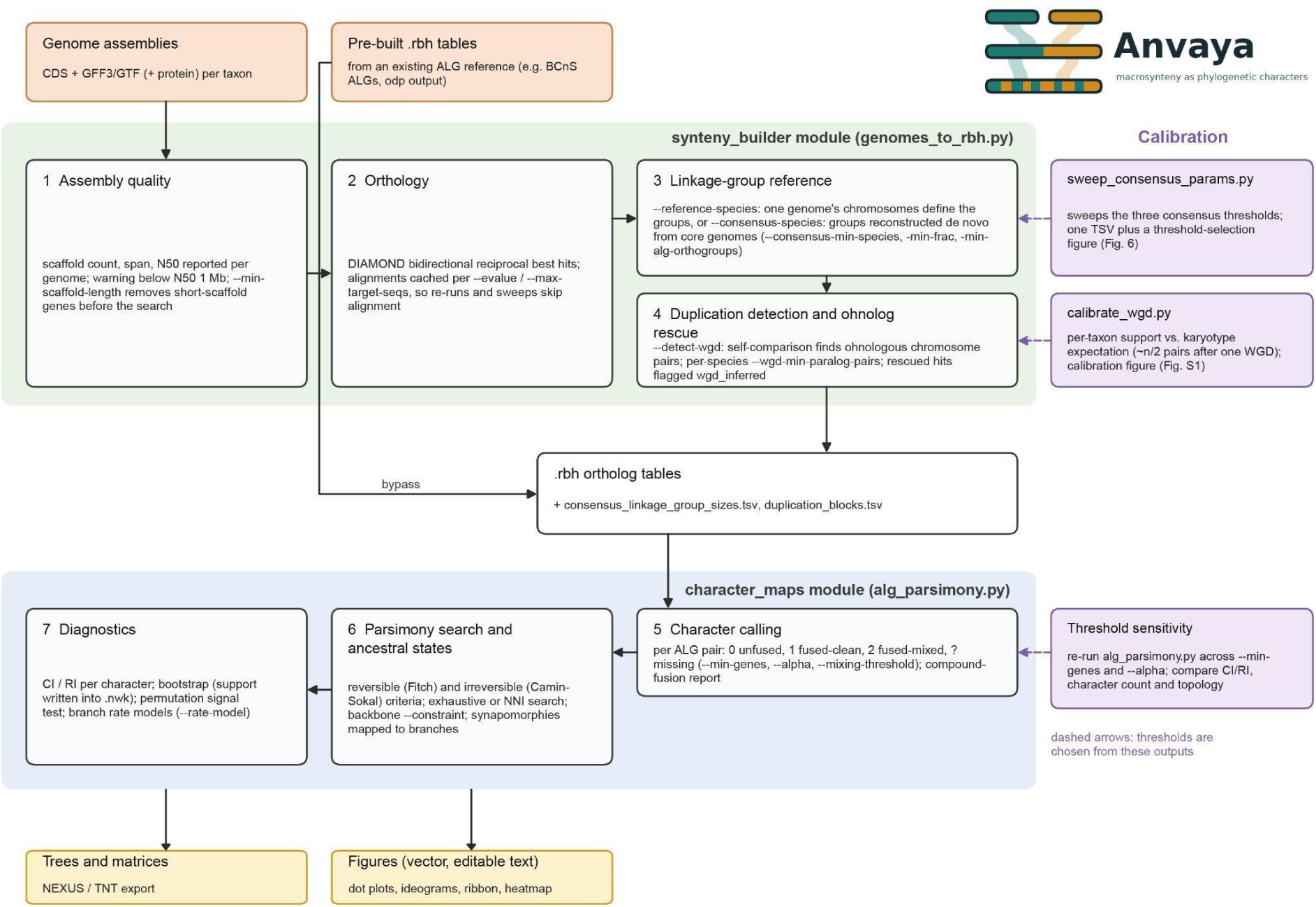
Anvaya workflow. The synteny_builder module (genomes_to_rbh.py; green) reports assembly contiguity (1), infers orthology by bidirectional reciprocal best hit with cached DIAMOND alignments (2), defines linkage groups from a single reference genome or by multi-genome consensus (3), and optionally detects duplication and rescues ohnologs (4), writing .rbh ortholog tables with linkage-group size and duplication-block reports. Pre-built .rbh tables, for example against the BCnS linkage groups or from odp, can bypass this module. The character_maps module (alg_parsimony.py; blue) codes each pair of linkage groups as unfused, fused-clean, fused-and-mixed or missing (5); searches for most-parsimonious trees under reversible and irreversible criteria, optionally under a backbone constraint, and maps synapomorphies (6); and computes homoplasy, support, signal and rate diagnostics (7). Outputs (yellow) are trees and matrices, with NEXUS and TNT export, and vector figures. Companion tools (purple) produce the outputs from which thresholds are chosen: sweep_consensus_params.py (Fig. 2), calibrate_wgd.py (Fig. S1) and threshold-sensitivity reruns of alg_parsimony.py (Fig. 3; Table S4). Dashed arrows show where those choices return to the pipeline.

#### Input data, assembly quality and orthology

Each input genome is supplied as coding sequences (CDS) with a GFF3 or GTF annotation, optionally with protein sequences. Before any search, genomes_to_rbh.py reports scaffold count, assembly span, longest scaffold and N50 for every genome with a warning where N50 falls below 1 Mb. Macrosynteny inference is acutely sensitive to contiguity because co-occurrence is assessed per scaffold: a linkage group split across fragments both loses genuine fusions and manufactures spurious ones (Liu et al., 2018). No hard threshold is imposed, since the appropriate floor depends on karyotype and genome size, but a fragmented genome is flagged as a poor choice of reference or core genome even where it remains usable as a comparison taxon. An optional minimum scaffold length (--min-scaffold-length) removes genes on short scaffolds from the protein set before the search rather than afterwards, so that a gene on a small fragment cannot occupy a reciprocal-best-hit slot that a genuine ortholog would otherwise fill. Scaffold lengths are read from GFF3 sequence-region pragmas where present and otherwise approximated, as a lower bound, from the furthest annotated feature.

Orthology between two proteomes is established by bidirectional reciprocal best hit using DIAMOND (Buchfink et al., 2021). Furthermore, alignments are cached against their search parameters, so that repeated runs over the same genomes, including every step of a threshold sweep, skip the alignment entirely, serving as a checkpoint. Gene coordinates are resolved from the annotation by matching several identifier attributes with isoform suffixes stripped, since annotation files differ in how they link coding sequences to genes. In the resulting .rbh table each row is one ortholog, with its linkage-group identity and the taxon’s own scaffold and coordinate.

#### Establishing the linkage-group reference

Linkage groups are defined in one of two ways: single-reference and consensus mode. In single-reference mode, one genome’s chromosomes serve directly as the linkage groups. The mode is fast, but its results are only as good as that genome’s contiguity, karyotype and its evolutionary proximity to the ancestral karyotype; a rearranged derived reference will manufacture spurious fusion calls. The reference is not itself a taxon in the analysis, just as an external set such as the Bilaterian–Cnidarian–Sponge (BCnS) linkage groups (Simakov et al., 2022) would not be.

In consensus mode, linkage groups are reconstructed *de novo* from a set of two or more core genomes. Orthogroups are built as connected components of the graph of reciprocal best hits among core genomes, and any component containing two genes from the same core genome is discarded as probably paralogous. An orthogroup must be present in a minimum number of core genomes (--consensus-min-species), and two orthogroups are placed in the same linkage group when they are co-located in at least a set fraction of the total core genomes containing both (--consensus-min-frac). Merging uses agglomerative clustering in which each proposed merge is re-validated against the clusters’ current majority placement, so that weak pairwise links cannot chain transitively into a single artefactual group. Groups with fewer member orthogroups than a set minimum are discarded (--consensus-min-alg-orthogroups), and the selected groups are labelled cALG1, cALG2 and so on in decreasing order of size. Non-core genomes are assigned to groups by majority vote of their reciprocal best hits against each core genome. Because the reconstruction depends only on the core set, every genome, core or not, is a taxon in the downstream analysis, and any analysis sharing the same core genomes and thresholds shares the same linkage groups.

These three thresholds interact, implicative to the linkage groups on which every later inference rests, yet none has a universally correct value. sweep_consensus_params.py runs the reconstruction across a grid of values and reports, for each combination, the number of linkage groups, their size distribution and the proportion of orthogroups retained and all three are plotted together (Fig. 2). Because the size threshold is a post-reconstruction filter, every value on that axis is derived from a single reconstruction, and because alignments are cached, a full sweep costs little more computation than one run. Only achievable agreement fractions are meaningful: with n core genomes the observed fractions are k/n, and a threshold set just above one of them silently behaves like the next. The software reports the k-of-n meaning of every value used, warns when a value sits just above one of these boundaries, and marks the selected combination on the sweep figure (Fig. 2).

**Figure 2.**
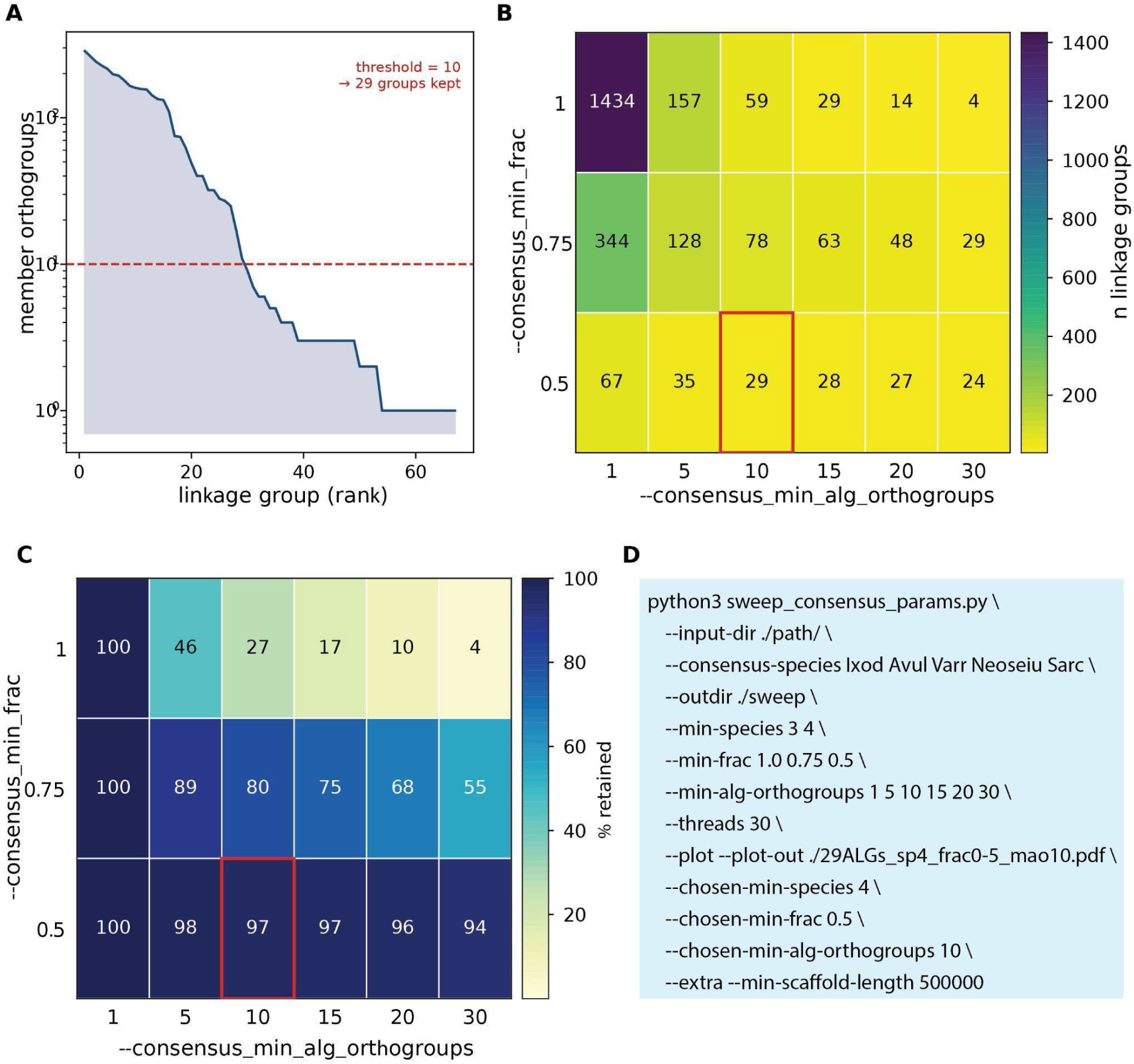
Choosing thresholds for consensus linkage-group reconstruction in the Acari dataset. Core genomes were *Ixodes scapularis*, *Argas vulgaris*, *Varroa destructor*, *Neoseiulus longispinosus* and *Sarcoptes scabiei*, with --consensus-min-species 4 and --min-scaffold-length 500000. (A) Reconstructed linkage groups ranked by number of member orthogroups at --consensus-min-frac 0.5. The dashed line marks the size threshold of ten orthogroups, which retains 29 groups; the distribution declines smoothly, so the threshold is a stated choice rather than a break in the data. (B) Number of linkage groups and (C) percentage of orthogroups retained across --consensus-min-frac (rows) and --consensus-min-alg-orthogroups (columns). Unanimous agreement (1.0) yields 1,434 candidate groups and retains as little as 4% of orthogroups once a size filter is applied; agreement of half the core genomes (0.5) retains 94–98% across all size thresholds. The chosen combination (0.5 and 10; 29 groups retaining 97% of orthogroups) is outlined in red. (D) The sweep_consensus_params.py command that produced panels A–C.

#### Whole-genome duplication detection and ohnolog rescue

Reciprocal best hit is one-to-one by construction, so when a genome carrying a lineage-specific duplication is compared with the reference, one member of every ohnologous pair is dropped (Catchen et al., 2009). The consequence is not a doubled signal but a false negative. Orthologs are split between the two retained copies of each ancestral chromosome, neither copy may reach the minimum needed to call an association, and a fully retained linkage group can vanish from the analysis.

For any species flagged as a possible duplication carrier (--detect-wgd), genomes_to_rbh.py compares the proteome with itself to find reciprocal-best-hit paralog pairs (Tang et al., 2008), excluding self-hits and pairs on the same scaffold, which are more likely tandem duplicates. Paralog pairs are aggregated by scaffold pair, and a pair of scaffolds is confirmed as ohnologous when it is supported by at least a set number of paralog pairs and passes a Fisher’s exact enrichment test against the genome-wide background of paralogy. Where one candidate pair is the only paralogy in the genome, the test has no background and the count threshold alone is applied. For each confirmed pair, every ortholog whose gene has a paralog on the partner scaffold gains that paralog as a second ortholog of the same reference gene, recovering the copy that reciprocal best hit discards. Every row records whether it came from a direct reciprocal best hit or from ohnolog rescue (flagged wgd_inferred), so the two kinds of evidence remain separable, and confirmed pairs are written to duplication_blocks.tsv. WGD rescue applies to the compared genome only; duplication in the reference genome, or in the core genomes that define the linkage groups, is not detected, which is why duplicated genomes should not be used to define linkage groups.

The minimum number of supporting paralog pairs can be set globally and overridden per species (for example, --wgd-min-paralog-pairs 30 Limulus=70), because paralog-pair counts scale with genome size, gene content and the number of duplication rounds, and a single global value would otherwise be dictated by the largest or most duplicated genome. Significance is a poor guide to this value. The enrichment test asks whether a chromosome pair carries more paralogs than the background predicts, which is not the same as asking whether the two chromosomes are ohnologous: two small chromosomes sharing a few paralogs can be significantly enriched. On empirical genome data, significance therefore accepts candidate pairs supported by a single paralog even after Bonferroni correction, and cannot be used to set the threshold. calibrate_wgd.py uses karyotype expectation instead. After one duplication each chromosome has one ohnologous partner, so roughly n/2 pairs are expected for n chromosomes, rising toward n where the genome has since been rearranged or duplicated again. For each taxon the tool reports the lowest thresholds consistent with each expectation and plots the support distribution, support against significance, and the numbers of confirmed pairs and of chromosomes with more than one partner across thresholds (Fig. S1). The last of these carries biological information of its own: chromosomes retaining several partners at every threshold indicate a network of ohnology rather than clean pairs.

#### Character construction: fusion and mixing

For each taxon, alg_parsimony.py first identifies which linkage groups are associated with which of its scaffolds. An association is called when it is supported by at least --min-genes orthologs and, where an enrichment p-value is available, when that value does not exceed --alpha. A scaffold associated with two or more linkage groups is a candidate fusion, and each pair on it is classified from gene order: genes are sorted by coordinate and the fraction of adjacent pairs that belong to different groups is computed. Above --mixing-threshold (default 0.3) the pair is coded fused-and-mixed (state 2); otherwise it is fused-clean (state 1), still arranged as two contiguous blocks. Pairs never co-located are coded unfused (0), and a pair is coded missing (?) in a taxon where either group has too few orthologs to assess, rather than being assumed unfused. The matrix contains one column for every pair of linkage groups fused in at least one taxon.

A scaffold carrying k linkage groups yields all C(k,2) pairwise characters, following convention (Simakov et al., 2022; Schultz et al., 2023). These characters are not independent, since they may record a single compound event, and treating them as independent inflates both the character count and apparent support. Anvaya therefore writes a report on every run that tabulates how many scaffolds carry each number of linkage groups and what share of all pairwise character instances each class generates, and warns when a minority of scaffolds accounts for most of them. In that case the effective number of independent fusion events is closer to the number of multi-group scaffolds than to the character count, and support values should be discounted accordingly.

#### Parsimony criteria and tree scoring

Trees are scored with a generalised Sankoff algorithm (Sankoff, 1975) parameterised by a cost matrix, so the two criteria differ only in the matrix supplied. Under the reversible criterion every change of state costs one, the unordered matrix equivalent to Fitch parsimony (Fitch, 1971). Under the irreversible criterion states are ordered (0 < 1 < 2), increases cost one and decreases are forbidden: fusion-with-mixing is treated as unable to revert (Schultz et al., 2023), in keeping with the rarity of fission and the persistence of fused domains observed across Lepidoptera (Wright et al., 2024). This is the Camin–Sokal model (Camin & Sokal, 1965) which differs from Dollo parsimony, which permits loss, and I chose it because the claim being modelled concerns irreversibility itself. The irreversible criterion requires a rooted tree; the root is placed on a specified outgroup, or on the taxon with the fewest derived states, which is flagged for the user to verify, and every character is fixed at state 0 at the root.

Because every change costs the same under the reversible criterion, and increases cost the same under the irreversible one, the reversible score of any tree can never exceed its irreversible score. This relationship is used as a consistency check on the search.

#### Tree search

For up to eight taxa by default (--max-exhaustive), every unrooted binary topology is enumerated by sequential taxon addition and scored under both criteria. Above that, the search is heuristic: random starting trees, built to satisfy any constraint, are improved by nearest-neighbour interchange until no further improvement is possible, and the best result over --nni-restarts starts is kept. After both searches, each criterion’s best trees are rescored under the other criterion. If a topology found under one criterion scores better under the other than that criterion’s own result, the search has stalled in a local optimum; the better topology is adopted and a warning is written to the report. The check cannot establish that a tree is optimal, only that it is not demonstrably suboptimal, so heuristic results should be confirmed with more restarts and in TNT or PAUP*.

#### Backbone constraints

A backbone constraint may be supplied as a Newick tree naming any subset of taxa, with any mixture of resolved and unresolved nodes. Each internal node asserts that its descendant taxa form a clade; a polytomy leaves relationships within that set unresolved. Candidate topologies are accepted only if every required clade is present once the topology is restricted to the taxa the constraint names, so that unnamed taxa may attach anywhere.

#### Ancestral states and synapomorphies

Ancestral states are reconstructed on the best tree under each criterion by Sankoff traceback (Clemente et al., 2009; Joy et al., 2016). States are assigned from the root downward so that each node minimises the combined cost of its subtree and the transition from its parent, with the root fixed at state 0 under the irreversible criterion. Every increase in state on an internal branch is reported as a synapomorphy of the clade above it; changes on terminal branches are counted in the score but not listed.

#### Homoplasy, support and signal diagnostics

Three diagnostics are reported with every tree. Consistency and retention indices (Kluge & Farris, 1969; Farris, 1989) are computed per character and for the whole matrix: CI = m/s and RI = (g − s)/(g − m), where m is the minimum possible number of steps, s the number on the reported tree and g the number on a completely unresolved tree, with missing data excluded. A character with CI below one has arisen more than once, and is not the single-origin synapomorphy it may appear to be. Clade support is estimated by nonparametric bootstrap, rerunning the full search on each replicate and writing support values into the Newick output. Clades implied by the choice of root alone are excluded. A permutation test asks whether the data carry phylogenetic signal at all: states are shuffled among taxa within each character, preserving each character’s composition and missing data, and the best achievable score is recomputed. The reported p-value is the fraction of replicates at least as parsimonious as the real data, with a +1/+1 correction, so that with R replicates the smallest attainable value is 1/(R + 1).

#### Visualisation

Six kinds of figures are drawn: Oxford dot plots, chromosome-painting ideograms, ribbon plots, character-matrix heatmaps, cladograms with synapomorphies on their branches, and duplication views in vector graphics with editable text. In dot plots each ortholog is placed by its position in its linkage group and in the taxon’s own scaffold; ideograms paint each scaffold by the linkage group of each successive ortholog; ribbon plots stack one painted track per taxon and join orthologs between adjacent tracks.

Several choices make ribbon plots of real data legible. Track order may follow a ladderised traversal of the inferred tree, so that clades are contiguous and crossings reflect rearrangement rather than listing order. Chromosome order within tracks may be optimised to minimise crossings using the barycenter heuristic of layered graph drawing, with the reduction in crossings reported. The linkage-group reference is stacked as the top track, where it serves as an unfused baseline; it is held fixed during optimisation because its order is meaningful. Gaps between chromosomes are set as a fraction of track width, with the total capped so that tracks with many chromosomes remain readable. Consecutive genes of one linkage group are drawn as a single segment, and runs of orthologs consecutive in both tracks as a single block ribbon with unique color assignment.

By default every recovered ortholog is drawn. With --plot-called-only, figures show only orthologs whose association is that pass the --min-genes and --alpha filters, so that a change of threshold is visible in the figures as well as in the tree. Where ohnologs have been rescued, rescued orthologs are marked in dot plots and ribbon plots, and a ribbon plot of each duplicated taxon against the reference alone shows its retained ohnologous chromosomes directly beneath the ancestral units.

#### Interoperability

The matrix can be exported as a NEXUS file with a PAUP* command block that searches under both unordered and irrev.up character types, PAUP*’s equivalents of the two criteria, honouring any backbone constraint (Swofford, 2003); and as a Hennig86 file with a TNT script (Goloboff et al., 2008). Either program can be run automatically if installed, but neither is required.

#### Empirical datasets and analysis settings

Two datasets were analysed (Table 1). The Acari dataset comprised thirteen chromosome-level genomes: seven Ixodida (two argasids and five ixodids), three Mesostigmata, two Acariformes and the pycnogonid *Endeis spinosa* as outgroup (Papadopoulos et al., 2025). The duplication analyses used the horseshoe crab *Limulus polyphemus*, whose lineage has undergone repeated rounds of duplication (Kenny et al., 2016; Nong et al., 2021), and the vinegaroon *Mastigoproctus giganteus*, which carries the single arachnopulmonate duplication (Kulkarni et al., 2024), with the unduplicated tick *Ixodes scapularis* as a baseline; the mesothele spider *Luthela* was included in threshold calibration only (Fig. S1). Consensus linkage groups were reconstructed in both datasets from the same five non-duplicated core genomes, one ixodid, one argasid, two mesostigmatans and one acariform (*Ixodes scapularis*, *Argas vulgaris*, *Varroa destructor*, *Neoseiulus longispinosus* and *Sarcoptes scabiei*), with --consensus-min-species 4, --consensus-min-frac 0.5, --consensus-min-alg-orthogroups 10 and --min-scaffold-length 500000, chosen from a sweep across --consensus-min-species 3–4, --consensus-min-frac 1.0, 0.75 and 0.5, and --consensus-min-alg-orthogroups 1–30 (Fig. 2). Because reconstruction depends only on the core set, both datasets share the same 29 linkage groups.

**Table 1.**
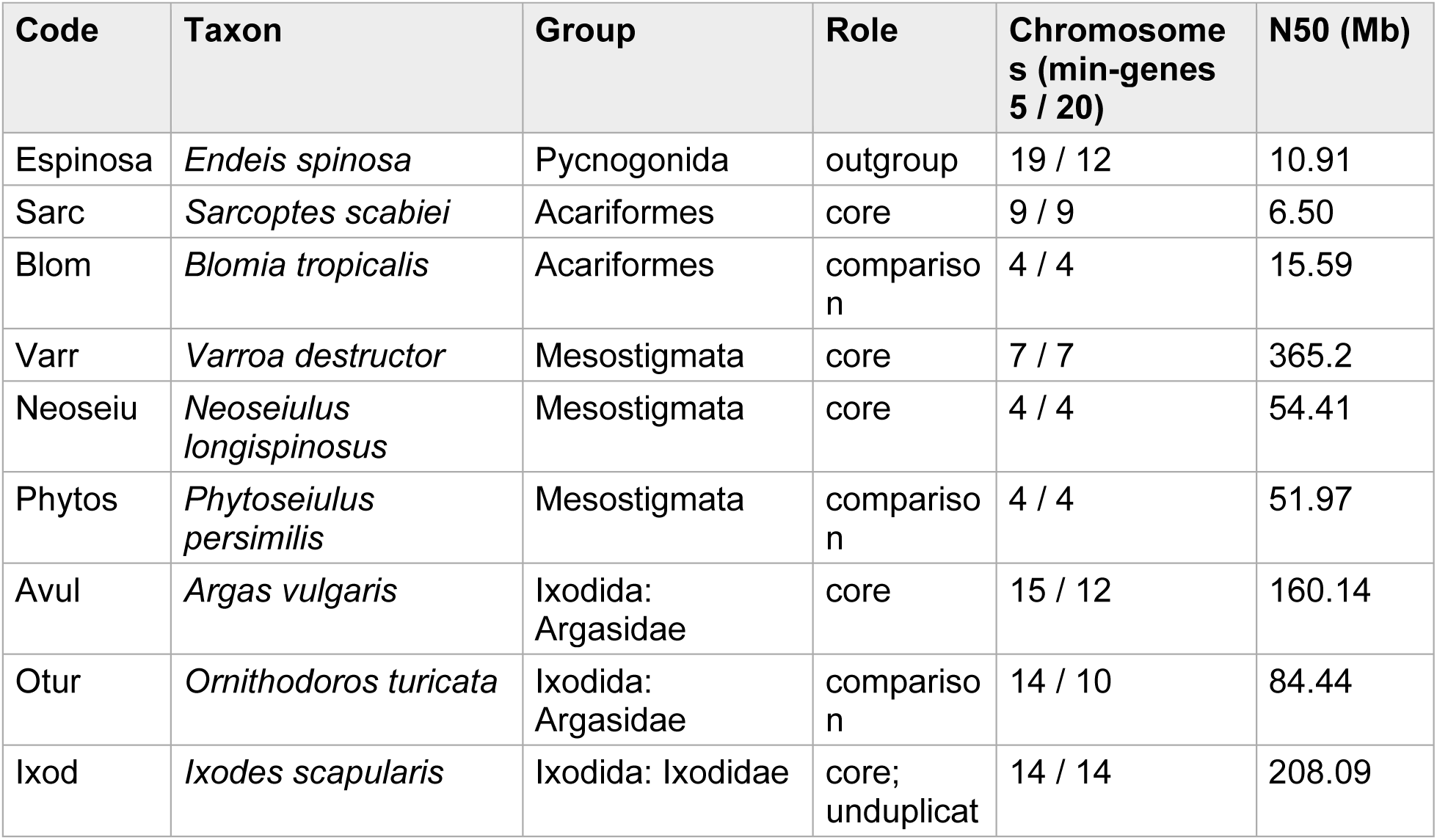
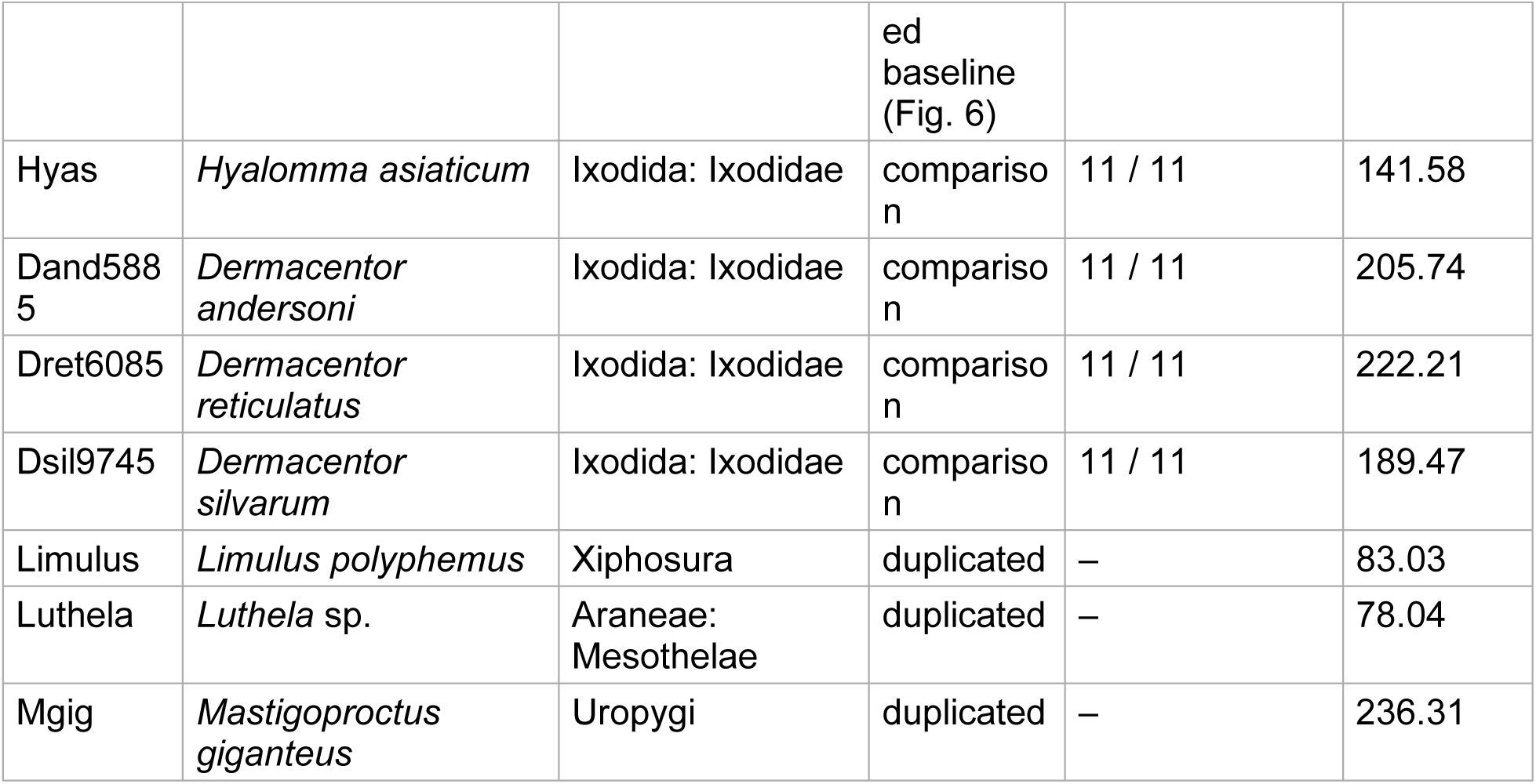
Genomes analysed. Chromosomes are those carrying at least one called linkage-group association at --min-genes 5 and 20 (Fig. 3). Core genomes defined the consensus linkage groups.

Trees were inferred with the pycnogonid as outgroup and with 1,000 bootstrap and 1,000 permutation replicates, under a backbone constraint that enforced the monophyly of Acariformes, Parasitiformes, Mesostigmata, Ixodida, Argasidae and Ixodidae and left relationships within Ixodidae and within Mesostigmata unresolved (Supplementary File S1). Characters were called at --alpha 0.05. Sensitivity to calling stringency was assessed by running the analysis at --min-genes 5 and 20 (Fig. 3), and by repeating it across --min-genes 5, 10 and 20 and --alpha 0.05 and 0.01 (Table S4); trees are reported for --min-genes 20.

**Figure 3.**
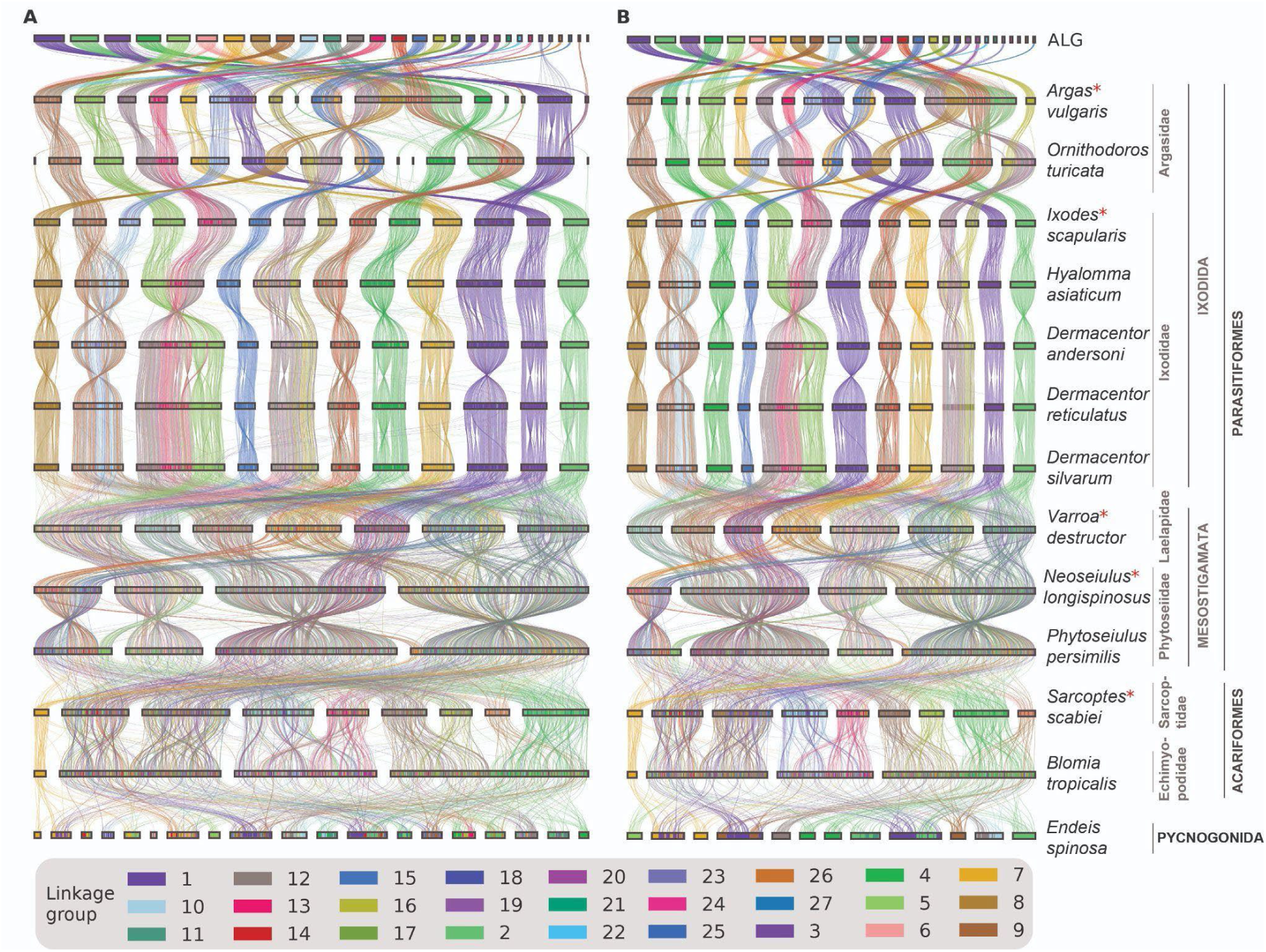
Sensitivity of macrosynteny calls to the ortholog minimum. Ribbon plots between the consensus linkage groups (top track) and thirteen genomes at (A) --min-genes 5 and (B) --min-genes 20, each drawing only associations that clear the respective threshold (--plot-called-only). Each track is one genome, painted by the linkage group of each ortholog; the top track is unfused by construction, and asterisks mark the five core genomes from which the linkage groups were reconstructed. Tracks are ordered by the constrained topology (Fig. 4B), with families and higher taxa indicated at right, and are rescaled to a common width, so horizontal extent reflects gene content rather than physical length; chromosome order within tracks minimizes ribbon crossings. Linkage group *n* in the figure corresponds to cALG*n* in the text. Two linkage groups present in (A) are absent from (B). The stricter threshold removes seven of the 19 chromosomes of *Endeis spinosa* with called associations, and unplaced scaffolds in the argasids, while leaving the ixodids, mesostigmatans and acariforms unchanged.

For the duplication analyses, candidate ohnologous pairs were first enumerated with the count filter disabled (--wgd-min-paralog-pairs 1), thresholds were chosen per taxon with calibrate_wgd.py, and .rbh tables were rebuilt with and without rescue for comparison. Figures were drawn with --plot-called-only.

## EMPIRICAL APPLICATION

Anvaya was applied to two questions in chelicerate genome evolution: the interrelationships of Acari, and the detection of whole-genome duplication in genomes with different duplication histories. The aim is to show how the software behaves on data of realistic size, and what it reveals about those data, rather than to offer a definitive phylogenetic result, which the limited taxon sampling would not support.

### Linkage-group reconstruction

The sizes of the reconstructed linkage groups decline smoothly, without a gap separating well-supported groups from fragments (Fig. 2A). The size threshold is therefore a stated choice rather than a property of the data. The agreement threshold matters far more (Fig. 2B, C). Requiring unanimity among core genomes fragments the reconstruction into 1,434 candidate groups; a size filter brings this down only by discarding most of the data, leaving 29 groups that retain 17% of orthogroups at a threshold of 15. Relaxing agreement to half of the core genomes gives 67 candidate groups, of which 29 have at least ten member orthogroups, and these retain 97% of all orthogroups. Retention varies little across size thresholds in this setting, from 98% at five orthogroups to 94% at thirty, so conclusions do not depend closely on the size cut-off chosen. The 29 groups are comparable in number to the 27 Acari-specific ancestral linkage groups defined independently for this clade (Bhoi et al., 2026).

### Macrosynteny across Acari and the effect of calling stringency

Ribbon plots show rearrangement that differs sharply among lineages (Fig. 3). The ixodid ticks retain comparatively coherent correspondence to the reconstructed units across eleven to fourteen chromosomes. This correspondence is expected in *Ixodes scapularis*, a core genome, but extends to the four metastriate ixodids, which did not contribute to the reconstruction. The four chromosomes of *Neoseiulus*, *Phytoseiulus* and *Blomia*, by contrast, each carry many linkage groups, the pattern produced by repeated fusion with mixing.

Raising the minimum from five (Fig. 3A) to twenty (Fig. 3B) supporting orthologs (--min-genes) removes two linkage groups entirely, because no taxon carries enough of their orthologs to call them. The effect is concentrated where calls are weakest. In the pycnogonid outgroup the number of chromosomes with a called association falls from 19 to 12, and in the argasids from 15 to 12 and from 14 to 10. The argasid losses are mostly unplaced scaffolds, and the ixodid, mesostigmatan and acariform genomes are unaffected. Associations supported by few orthologs therefore concentrate in the most distant genome and in the least contiguous parts of assemblies, which is precisely where a spurious fusion call would do most harm to the character matrix.

### Fusion-and-mixing characters and trees

At --min-genes 20, the matrix comprised 250 characters (Fig. 4A). Derived states are concentrated in the three mesostigmatans and in *Blomia tropicalis*, most of them fused-and-mixed, whereas *Sarcoptes scabiei* carries comparatively few; ticks carry fewer derived states, predominantly fused-clean, many shared across Ixodida. The pycnogonid outgroup is almost entirely unfused. Most reconstructed transitions are direct changes from unfused to fused-and-mixed. Only two proceed from a clean fusion to mixing within the tree (cALG12+cALG19 and cALG14+cALG6), each preceded by a clean fusion earlier in the tree, which is the ordered sequence the three-state coding is designed to capture.

**Figure 4.**
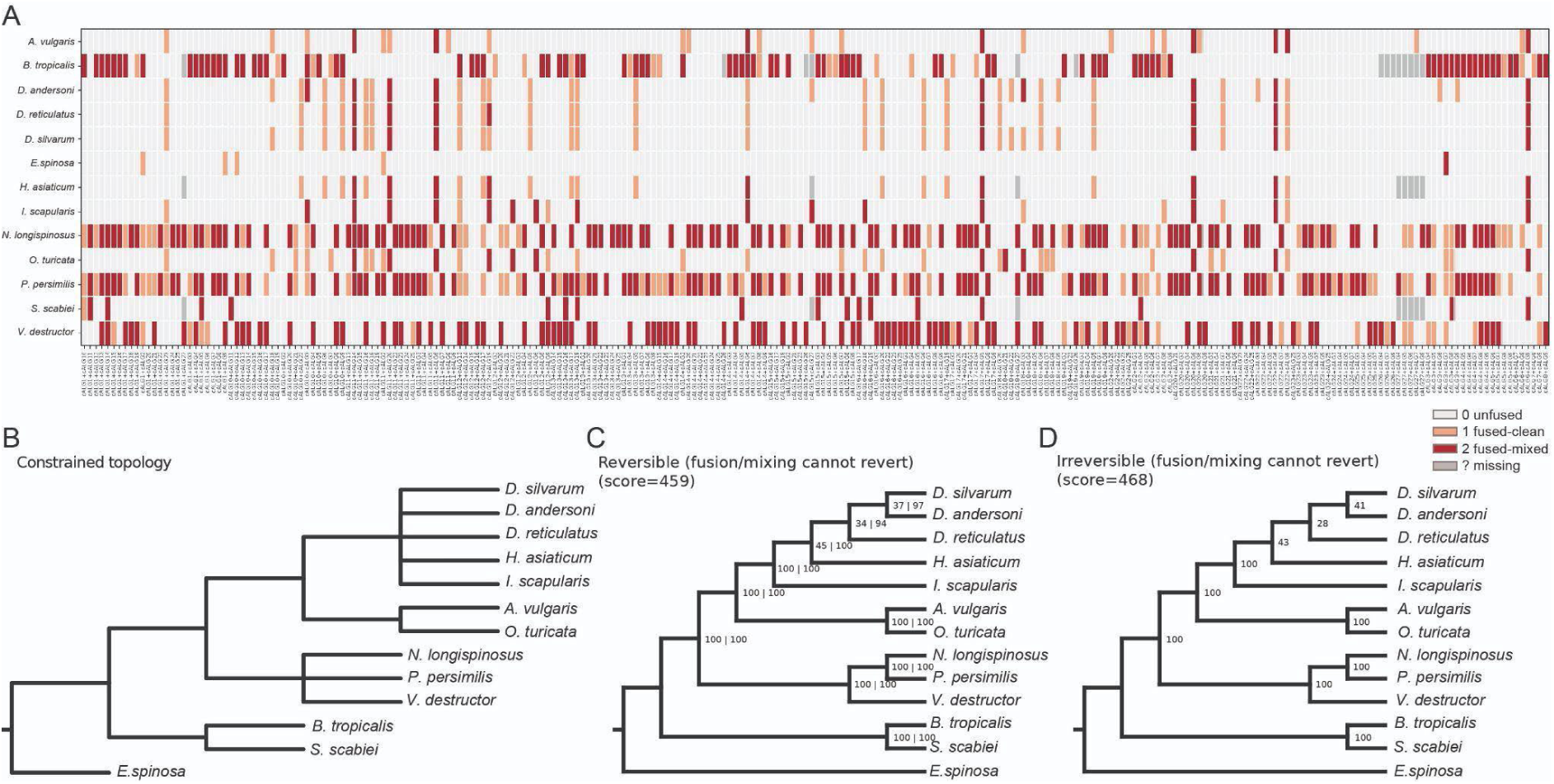
Fusion-and-mixing characters and the resulting trees for the Acari dataset at --min-genes 20. (A) Taxa-by-characters matrix; each column is one pair of linkage groups fused in at least one taxon, coded unfused (0), fused-clean (1), fused-and-mixed (2) or missing (?); 250 characters, 13 taxa. (B) Backbone constraint applied in both searches, enforcing Acariformes, Parasitiformes, Mesostigmata, Ixodida, Argasidae and Ixodidae and leaving relationships within Ixodidae and within Mesostigmata unresolved. (C) Most-parsimonious tree under the reversible criterion (score 459) and (D) under the irreversible criterion, in which fusion and mixing cannot revert (score 468), both rooted on *Endeis spinosa*. The two criteria recover the same topology. Node values are bootstrap support from 1,000 replicates with Anvaya/TNT version 1.5 respectively; support for constrained clades is 100% by construction and is not evidence for them. Ensemble CI 0.754, RI 0.742; permutation test p < 0.001 (1,000 replicates).

Both criteria recovered the same topology (Fig. 4C, D), scoring 459 steps under the reversible criterion and 468 under the irreversible criterion; the nine-step difference being the cost, on this topology, of the imposed irreversibility rule. An initial heuristic search had returned the opposite ordering, a reversible tree scoring 479 against 478 for the irreversible tree. That pair is impossible for optimal trees, since the reversible score of any topology cannot exceed its irreversible score, and showed that the reversible search had stalled in a local optimum. This observation prompted the cross-check (see Tree search section), and the trees reported here were re-estimated with it and --nni-restarts 500. The backbone constraint enforced the established monophyly for Acariformes, Parasitiformes, Mesostigmata, Ixodida, Argasidae and Ixodidae (Fig. 4B). Their 100% bootstrap support follows from searching every replicate under the same constraint and so is expected. The relationships left free were those within Ixodidae and within Mesostigmata. Both criteria placed *Ixodes scapularis* sister to the remaining ixodids, consistent with the established division between Prostriata and Metastriata; recovered *Dermacentor* as monophyletic, with *D. silvarum* and *D. andersoni* as sister species; and grouped the two phytoseiids, *Neoseiulus* and *Phytoseiulus*, to the exclusion of *Varroa*. The phytoseiid pair received full bootstrap support under both criteria, whereas the three metastriate nodes were weakly supported (34–45% under the reversible criterion and 28–43% under the irreversible criterion). Ensemble CI and RI were 0.733 and 0.712. The permutation test gave p < 0.001 (1,000 replicates; the smallest attainable value), so the matrix carries more signal than randomly structured data of the same composition.

These figures must be read together with two further results. Across --min-genes 5, 10 and 20 (Table S4), the character count fell by 29.8% (356→250) while ensemble CI rose from 0.498 to 0.754 under the irreversible criterion, suggesting that associations called on few orthologs add homoplasy rather than information; --alpha had comparatively little effect. The compound-fusion report showed that 34.5% of scaffolds carrying five or more linkage groups generated 86.6% of all pairwise character instances, so the number of independent fusion events is well below the number of characters. Macrosynteny here complements sequence data rather than replacing it, and a large character count should not be read as strong evidence, consistent with recent assessments of its limits in arachnids (Copley, 2025; Bhoi et al., 2026; Kulkarni et al., 2026).

### Calibrating duplication detection

In all three duplicated genomes, significance alone failed to identify ohnologous chromosome pairs (Fig. S1). After Bonferroni correction for the number of chromosome pairs tested, 154 of 325 candidate pairs in *Limulus* (47%) and 55 of 153 in *Mastigoproctus* (36%) were significant, as were pairs supported by a single paralog in *Limulus*, three in *Mastigoproctus* and seven in *Luthela*. Were these accepted, nearly half of all possible chromosome pairs in *Limulus* would be ohnologous, which is impossible. Support and significance are only loosely related in all three genomes (Fig. S1, B1–B3).

Thresholds chosen from karyotype expectation (70, 15 and 30 supporting paralog pairs for *Limulus*, *Luthela* and *Mastigoproctus*) retained 13, 24 and 8 ohnologous pairs, against roughly 13, 25 and 9 expected after a single duplication of 26, 50 and 18 chromosomes. The thresholds differ almost five-fold, yet they apply one criterion consistently, and at each a similar minority of chromosomes (23%, 30% and 22%) retains more than one partner. *Mastigoproctus* alone shows a genuine break in support: the number of confirmed pairs is stable at about 70 up to a threshold of ten, then falls steeply to eight pairs at thirty, where its threshold lies. For *Limulus* the single-duplication expectation is conservative, because more than one round of duplication is expected to leave chromosomes with several partners (Kenny et al., 2016; Nong et al., 2021), and a threshold of 37 would retain the 26 pairs of the permissive expectation. I kept the conservative value, so rescued counts for *Limulus* are lower bounds.

### Ohnolog rescue in three duplicated genomes

Ohnolog rescue recovered 205 of 1,636 orthologs in *Limulus polyphemus* (12.5%) and 44 of 1,566 in *Mastigoproctus giganteus* (2.8%), and its consequences differed accordingly (Fig. 5). Restricting the figures to associations that cleared the ortholog minimum (--plot-called-only) makes the effect explicit. In *Limulus*, 20 chromosomes carried a called linkage group without rescue and 23 with it (Fig. 5A, B): Chr11, Chr23 and Chr24 appear only after rescue. Without it their orthologs were divided between the two retained copies of each ancestral chromosome, neither copy reached the minimum of twenty, and all three chromosomes were called as carrying no ancestral linkage group — the false negative (described in Whole-genome duplication detection and ohnolog rescue section) in which a retained duplication is rendered invisible rather than conspicuous. In *Mastigoproctus* the same 14 scaffolds were called with and without rescue (Fig. 5C, D); the rescued orthologs reinforced existing associations but created none, as expected where an ancient duplication has since been eroded by gene loss and rearrangement. In neither genome did rescue change the number of linkage groups called (15 and 16 of 29 respectively); it recovers chromosomes, not groups. The contrast between the two genomes is informative in itself. Rescue matters most where duplicates are well retained and the two copies are similar enough that reciprocal best hit must choose between them, and least where divergence and loss have already reduced one copy below detection. The magnitude of the effect, 12.5% against 2.8% of orthologs, is therefore a rough index of how much of a duplication survives in a comparable state, and should be reported alongside the number of confirmed ohnologous chromosome pairs rather than in place of it.

**Figure 5.**
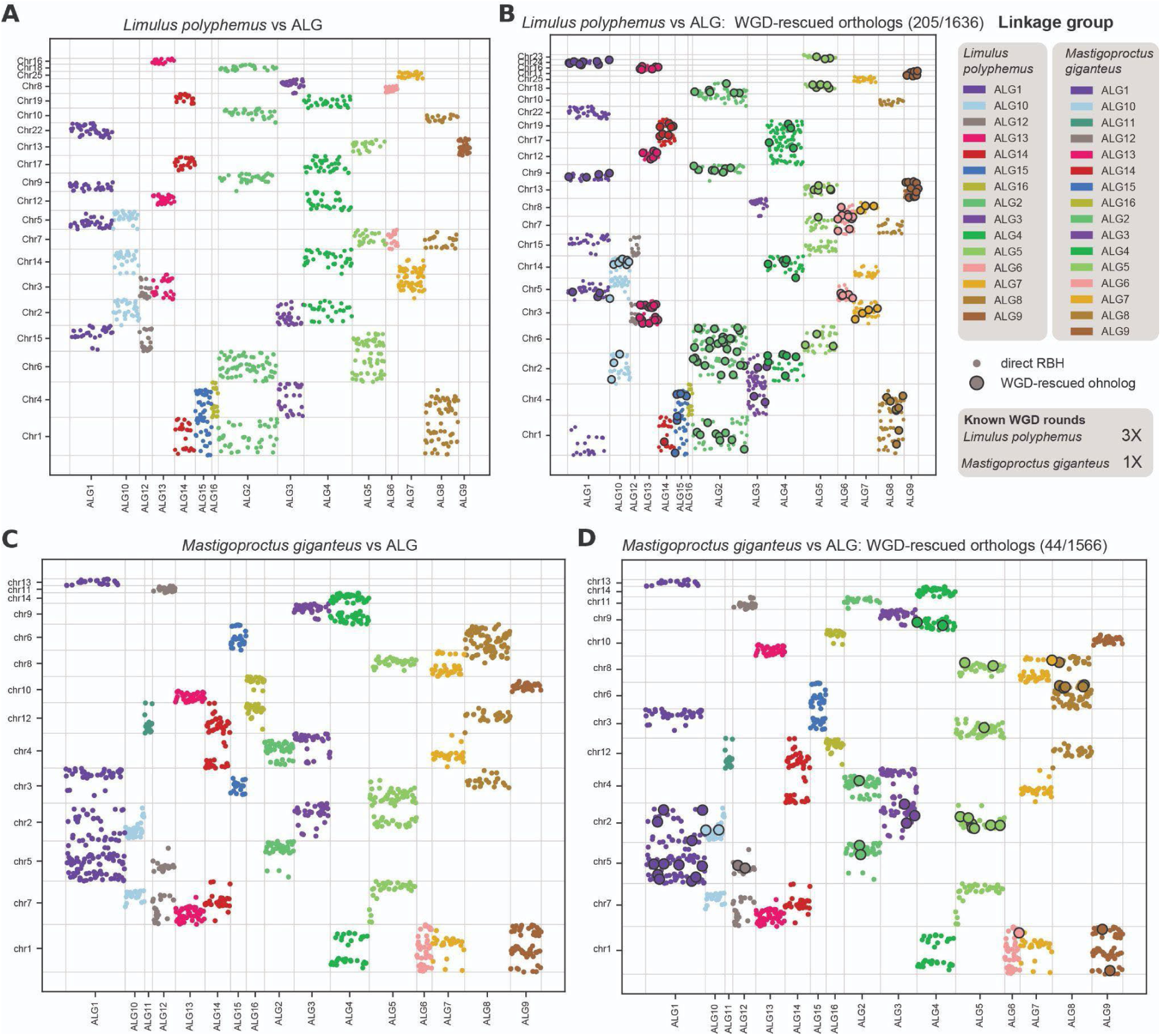
Effect of ohnolog rescue in two duplicated genomes. Oxford dot plots of (A, B) *Limulus polyphemus* and (C, D) *Mastigoproctus giganteus* against the consensus linkage groups, (A, C) without and (B, D) with --detect-wgd. Each point is one ortholog, placed by its position within its linkage group (X-axis) and within the taxon’s own chromosome or scaffold (Y-axis), and coloured by linkage group. All panels show only associations supported by at least 20 orthologs (--plot-called-only), which leaves 15 linkage groups in *Limulus* and 16 in *Mastigoproctus*; in B and D, orthologs recovered by rescue are enlarged and ringed. Rescue recovered 205 of 1,636 orthologs in *Limulus* and 44 of 1,566 in *Mastigoproctus*. In *Limulus*, 20 chromosomes carry a called association without rescue and 23 with it, Chr11, Chr23 and Chr24 appearing only after rescue; in *Mastigoproctus*, the same 14 scaffolds are called either way. The inset gives the known number of whole-genome duplication rounds (three in *Limulus*, one in *Mastigoproctus*). Chromosomes are ordered by ortholog count, so row order differs between panels. Thresholds: --wgd-min-paralog-pairs Limulus=70, Mgig=30.

Ribbon plots against the reference show the copy number of ancestral units directly, with the unduplicated tick *Ixodes scapularis* as a baseline (Fig. 6). In *Ixodes* each linkage group maps predominantly to a single one of its 14 chromosomes, several of which carry more than one group (Fig. 6A); because *Ixodes* is a core genome, this close correspondence is partly a consequence of construction. In *Limulus*, by contrast, single linkage groups are distributed across up to six chromosomes; cALG1, for example, occurs on Chr1, Chr5, Chr9, Chr15, Chr22 and Chr24, consistent with repeated rounds of duplication (Fig. 6B). In *Mastigoproctus*, groups occupy up to four scaffolds, as for cALG1 and cALG5, consistent with a single ancient duplication followed by fission and fusion (Fig. 6C; Kulkarni et al., 2024).

**Figure 6.**
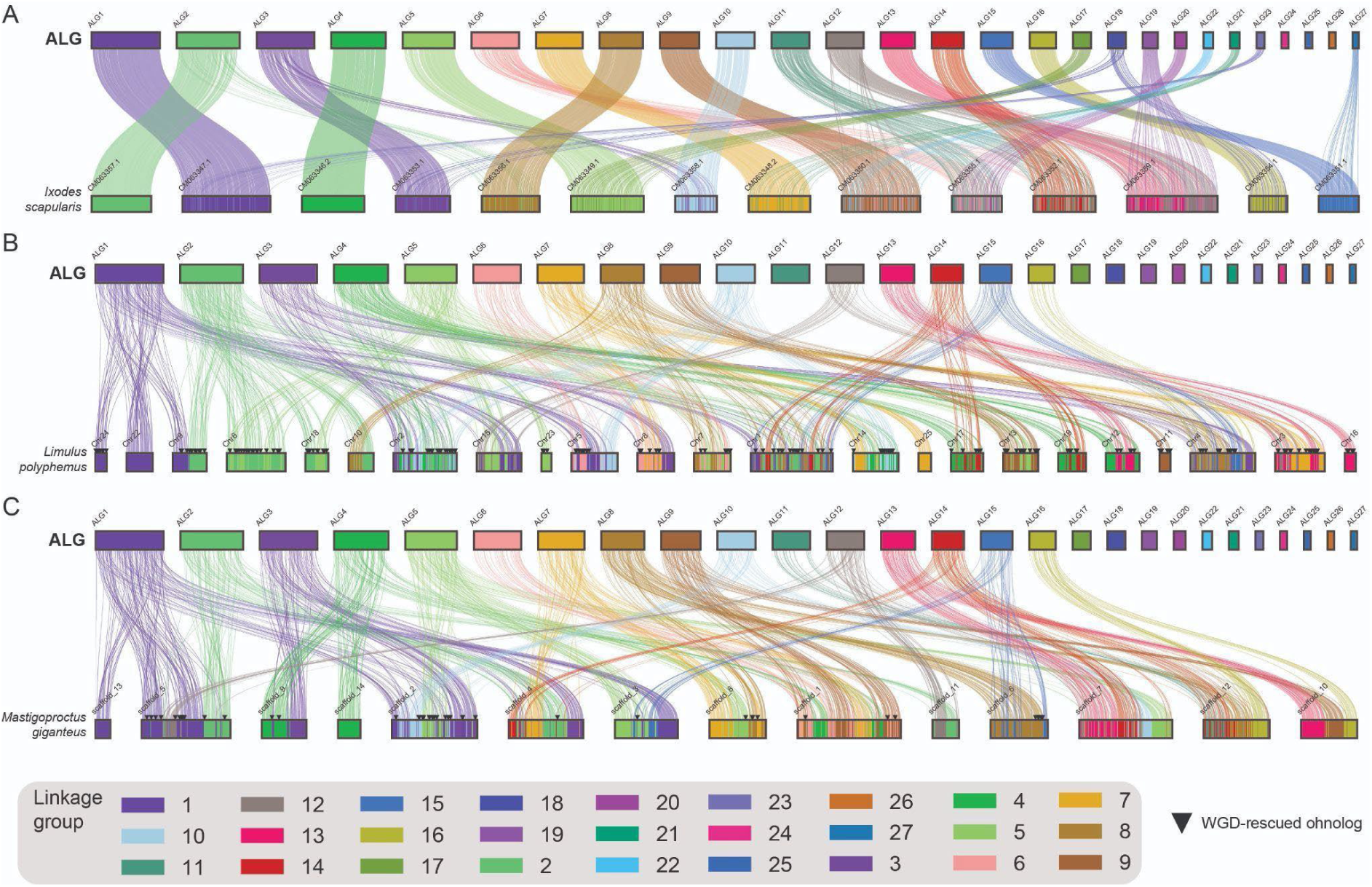
Consensus linkage groups in an unduplicated genome and two duplicated genomes. Ribbon plots of the consensus linkage groups (top track) against (A) the tick *Ixodes scapularis*, which has no known whole-genome duplication, (B) the horseshoe crab *Limulus polyphemus* and (C) the vinegaroon *Mastigoproctus giganteus*, each drawn against the reference alone; triangles mark rescued ohnologs. In *Ixodes* each linkage group maps predominantly to a single chromosome, several chromosomes carrying more than one group; because *Ixodes* was a core genome, this correspondence is close partly by linkage group design. Ancestral units occupy up to six chromosomes in *Limulus*, consistent with more than one round of duplication, and up to four scaffolds in *Mastigoproctus*, consistent with a single duplication followed by rearrangement. Only associations supported by at least 20 orthologs are drawn (--plot-called-only).

Only 15 of the 29 linkage groups in *Limulus*, and 16 in *Mastigoproctus*, reached twenty supporting orthologs. The smaller consensus groups, cALG17–29 in both genomes and cALG11 in *Limulus*, contribute too few orthologs in these distant genomes to be called at this stringency. Duplication analyses of distant taxa therefore rest on the larger linkage groups, and a lower --min-genes, applied to these taxa only, would bring the smaller ones into view at the cost of weaker individual calls.

## Discussion

### Relationship to existing softwares

Anvaya fills a gap between existing tools rather than replacing any of them. odp remains the standard for reciprocal-best-hit orthology against a reference set, enrichment testing and Oxford plots (Schultz et al., 2023), and Anvaya follows its conventions. macrosyntR tests chromosome associations pairwise (El Hilali & Copley, 2023). Syngraph (Mackintosh et al., 2023) and LINGUINE (Vargas-Chávez & Fernández, 2026) reconstruct ancestral linkage groups at the nodes of a given tree, and egt measures continuous mixing and rates of change along a calibrated tree (Schultz & Simakov, 2026). Each of these either compares genomes pairwise or takes the tree as given. Anvaya instead infers the tree, treating fusion and mixing as discrete characters under stated cost models, with output checkable in TNT or PAUP*.

LINGUINE is the closest recent framework and differs in two further respects. It builds orthogroups or hierarchical orthogroups and delineates syntenic blocks with a hidden Markov model, gaining markers and tolerance of gene loss and fragmentation; Anvaya’s reciprocal best hit is simpler and faster but recovers fewer markers and handles many-to-many orthology less well. Where a trusted tree or curated orthogroups exist, LINGUINE’s reconstruction is better resourced, and its output can be supplied to Anvaya as a reference. The two also treat duplication differently, for different purposes. LINGUINE collapses duplicated linkage groups before reconstruction, which is appropriate when duplication confounds inference of ancestral states. Anvaya recovers the dropped ohnolog because the duplication is itself the signal of interest. Rescued orthologs are flagged throughout, so that they are never mistaken for independent direct evidence. I have not benchmarked the two directly, because their outputs, ancestral states on a fixed tree and a tree inferred from characters, are not commensurable; a fair comparison would need data with both a known tree and known ancestral states.

### What the empirical analyses show about the method

Three observations from the empirical analyses apply beyond these data. First, thresholds used for calibration have effects on the inference. Weak associations concentrate in the most distant genome and in unplaced scaffolds, so raising the ortholog minimum removes them preferentially (Fig. 3; Table S4), and the agreement threshold for consensus reconstruction changes the number of linkage groups by two orders of magnitude. Every threshold should be reported with a sensitivity analysis rather than as a single value, and Anvaya’s sweep and calibration tools exist to make that a general practice.

Second, significance is not evidence of ohnology. In all three duplicated genomes, Bonferroni-corrected enrichment accepted chromosome pairs supported by one to seven paralogs, and in *Limulus* nearly half of all possible pairs. Karyotype expectation is a more useful constraint, and the number of chromosomes that retain several partners is informative in its own right: it separates the multiple rounds of duplication in horseshoe crabs from the single, since-eroded ancient duplication of arachnopulmonates (Fig. S1; Fig. 6). Ohnolog rescue restores chromosomes that reciprocal best hit hides (Fig. 5). It matters most where duplicates are well retained, as in the xiphosuran *L. polyphemus*, and least where an ancient duplication has since decayed, as in the uropygid *M. giganteus*.

Third, heuristic searches must be checked. In an initial run of the empirical analysis, the reversible search returned a tree scoring worse than one already found under the irreversible criterion, which is impossible for optimal trees under these cost matrices. This relationship between the criteria is a free diagnostic that now runs automatically. It cannot suggest an optimum, however, trees from datasets should be confirmed with more restarts and in TNT or PAUP*.

### Why parsimony

Probabilistic models of karyotype evolution accommodate heterogeneous rates of rearrangement that parsimony does not (Yoshida & Kitano, 2021), and rate heterogeneity is real: fission and fusion rates differ among lineages of Lepidoptera (Wright et al., 2024), and macrosyntenic characters can recapitulate rather than resolve conflict among rapidly diversifying lineages (Kulkarni et al., 2026). I use parsimony for three reasons: (1) It requires no rate model, which matrices of this size could not estimate reliably; (2) It makes irreversibility an explicit, testable cost assumption, allowing the same data to be scored with and without it; (3) It maps synapomorphies onto branches in a form that can be read directly as cladistic evidence.

### Assumptions and limitations

Fusion is called from whole-scaffold association only, so breakpoint and fission signal within scaffolds is not modelled. Decomposing compound fusions into pairwise characters follows convention but violates the independence parsimony assumes; the effect is now measured and reported, but not yet corrected. Enrichment tests of the kind used here can overstate significance (Copley, 2025), which is one reason every threshold is exposed as a parameter. Single-reference mode inherits the reference genome’s own rearrangements and assembly errors. Consensus mode is more robust but depends on the core set, and, like the reference in single-reference mode, core genomes must not carry their own duplication. Orthogroups in consensus mode are connected components of a reciprocal-best-hit graph, which is simpler than dedicated orthology inference (Emms & Kelly, 2019) and may be misled by chains of hits in paralog-rich genomes; ambiguous components are discarded, but this is a coarse safeguard.

Ohnolog rescue assumes that duplication resolves into pairs of chromosomes. Where chromosomes have several partners, as in *Limulus*, or duplicates have diverged or been lost, as in *Mastigoproctus*, recovery is partial and rescued counts are lower bounds. Calibration against karyotype expectation requires a reliable chromosome count, which scaffold counts may not provide. The smaller consensus linkage groups may fall below the ortholog minimum in distant genomes and so contribute nothing to their analysis. When many trees tie, one representative is reported with the number of ties; consensus trees are not yet computed. The heuristic search is a nearest-neighbour hill-climber whose results should be confirmed externally, and bootstrap and permutation replicates share its limitations. Scaffold lengths inferred from annotation are lower bounds, so length filters should be set conservatively.

### Future directions

The most immediate extensions follow from these limitations: strict and majority-rule consensus of tied trees; Bremer support to complement the bootstrap; down-weighting of characters that arise from one compound fusion; duplication-aware orthology for consensus reconstruction; stronger tree search, including tree-bisection-reconnection; and rescue that accommodates several ohnologous partners per chromosome and duplication in the core genomes. None is needed for the present use of the software, but each would widen the conditions under which its results can be trusted.

### Software availability

Anvaya (genomes_to_rbh.py, alg_parsimony.py, sweep_consensus_params.py, calibrate_wgd.py and make_toy_dataset.py) is implemented in Python 3 and released at https://github.com/kulkarni-lab/anvaya. The synthetic validation datasets and the exact commands used to produce every figure and table are provided as supplementary material at [figshare link].

---

**Figure S1.** Per-taxon calibration of --wgd-min-paralog-pairs for *Limulus*, *Luthela* and *Mastigoproctus* (rows). (A) Candidate ohnologous chromosome pairs ranked by supporting paralog pairs, coloured by Bonferroni-corrected significance; the dashed line marks the chosen threshold (70, 15 and 30), which confirms 13, 24 and 8 pairs. (B) Supporting paralog pairs against significance, with the Bonferroni level for the number of pairs tested (1.5 × 10⁻⁴, 4.1 × 10⁻⁵ and 3.3 × 10⁻⁴); significant pairs occur at every level of support, so significance alone does not identify ohnologous chromosomes. (C) Numbers of confirmed pairs (blue) and of chromosomes with more than one partner (red) across thresholds; thresholds were chosen where confirmed pairs fall to the single-duplication expectation of about one partner per chromosome.

## ACKNOWLEDGEMENTS

All analyses were conducted on the Ramanujan High Performance Computing at the CSIR-CCMB, Hyderabad, India. I am grateful to Pratik Khopkar, Arjun Cherukutty and Nithin KA for their comments on the previous drafts. Funding for this study came from the Ramanujan Fellowship (ANRF/RJF/2023/000045, S.K.), Early Career Research Grant (ANRF/ECRG/2024/000947/LS, S.K.) and OLP0030 (S.K.) grants.

---

**Table S1.** Character states recovered for the five-genome synthetic dataset. Five genomes (G1–G5) were constructed with twelve orthologous genes descended from three ancestral linkage groups, four genes each, and character states assigned by design rather than evolved along a tree. G1 and G4 were built with all three linkage groups unfused; G2 and G5 with the first two fused as two contiguous blocks; G3 with the same two fused and their genes fully interleaved. Consensus linkage groups were reconstructed from G1, G4 and G2 as core genomes, deliberately including one genome (G2) in which the two groups are fused. For the single character distinguishing the first two linkage groups, Anvaya recovered state 0 (unfused) in G1 and G4, state 1 (fused-clean) in G2 and G5, and state 2 (fused-and-mixed) in G3 — all five as designed. Default thresholds (--min-genes 3, --alpha 0.05, --mixing-threshold 0.3).

**Table S2.** Effect of ohnolog rescue on the simulated duplicated genome. A non-duplicated genome (TICK) carrying two four-gene linkage groups on separate scaffolds was compared with a duplicated genome (SPIDER) carrying two ohnologous copies of the same eight-gene set. Which ohnolog won each reciprocal best hit was alternated by gene index, so that under plain RBH each linkage group is split two-and-two across the two ohnologous scaffolds rather than concentrated on one. Without --detect-wgd, 8 of 16 true orthologs were recovered and neither linkage group reached the minimum of three on any scaffold, so no linkage group was called anywhere in SPIDER. With --detect-wgd, the intragenomic self-comparison found 8 reciprocal-best-hit paralog pairs, all connecting the same two scaffolds; the pair was confirmed on paralog-pair count alone, since it was the only paralogy signal in this small genome and the enrichment test was therefore uninformative. All 16 orthologs were recovered and both linkage groups were correctly called on both ohnologous scaffolds (--min-genes 3, --wgd-min-paralog-pairs 5).

**Table S3.** Per-character consistency and retention indices for the six-taxon diagnostic dataset, identical under both criteria. The matrix encodes the known tree (T1,(T2,((T3,T4),(T5,T6)))) with four fusion characters: three clean, nested synapomorphies (A+B uniting T3–T6; C+D uniting T5 and T6; E+F uniting T3 and T4) and one deliberately homoplastic character (G+H, present in T2 and T5, which do not form a clade). m, minimum possible steps; s, steps on the recovered tree; g, steps on a fully unresolved tree. Each designed synapomorphy scores CI = RI = 1.000; the homoplastic character is correctly and specifically flagged at CI = 0.500, RI = 0.000. Ensemble CI = Σm/Σs = 4/5 = 0.800 and RI = (Σg − Σs)/(Σg − Σm) = (8 − 5)/(8 − 4) = 0.750, both matching the hand-computed expectation.

