## Supplemental Table for "Anvaya: phylogenomic analysis of chromosome fusion-and-mixing characters and detection of shared whole-genome duplication"

Supplementary figures

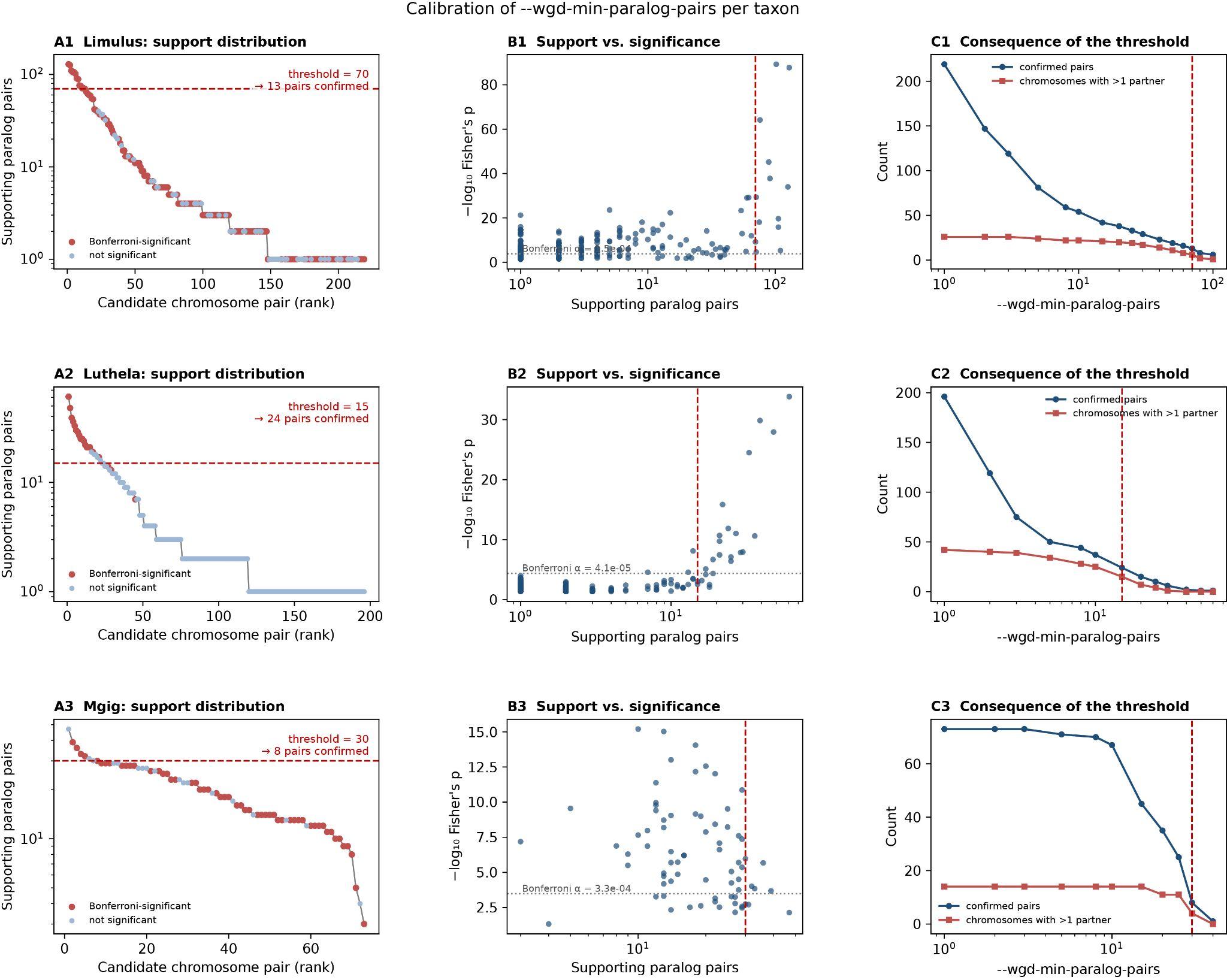

**Figure S1.** Per-taxon calibration of --wgd-min-paralog-pairs for *Limulus*, *Luthela* and *Mastigoproctus* (rows). (A) Candidate ohnologous chromosome pairs ranked by supporting paralog pairs, coloured by Bonferroni-corrected significance; the dashed line marks the chosen threshold (70, 15 and 30), which confirms 13, 24 and 8 pairs. (B) Supporting paralog pairs against significance, with the Bonferroni level for the number of pairs tested (1.5 × 10⁻⁴, 4.1 × 10⁻⁵ and 3.3 × 10⁻⁴); significant pairs occur at every level of support, so significance alone does not identify ohnologous chromosomes. (C) Numbers of confirmed pairs (blue) and of chromosomes with more than one partner (red) across thresholds; thresholds were chosen where confirmed pairs fall to the single-duplication expectation of about one partner per chromosome.

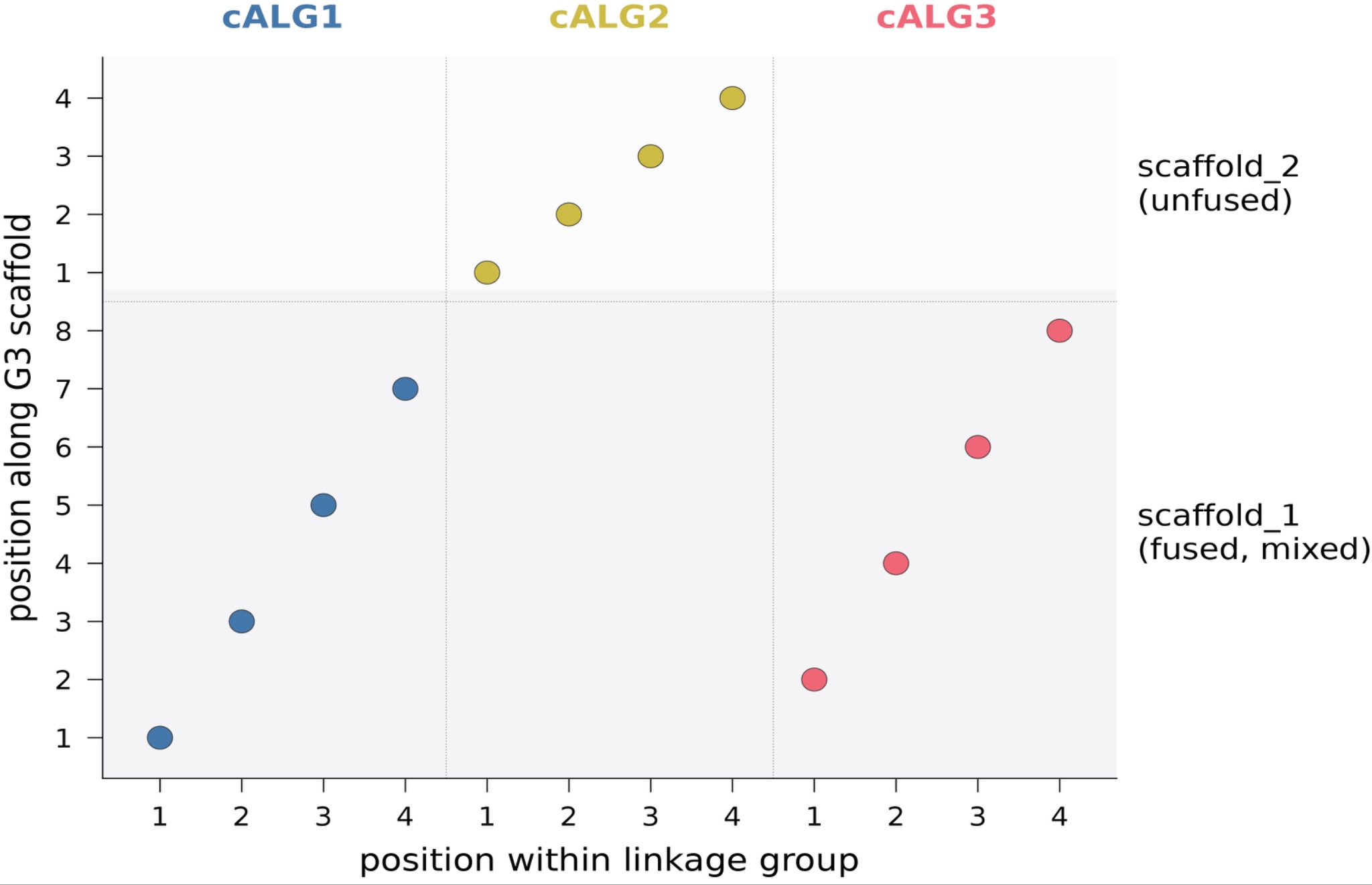

**Figure S2.** Oxford-style synteny dot plot for the simulated genome G3 against the reconstructed consensus linkage groups (x-axis) and G3's own scaffolds (y-axis). Genes of the two fused ancestral linkage groups (cALG1 and cALG3) are visibly interleaved along the shared scaffold, the signature that the mixing statistic is designed to detect, in contrast to the clean single-block diagonal of the unfused third group (cALG2) on its own scaffold.

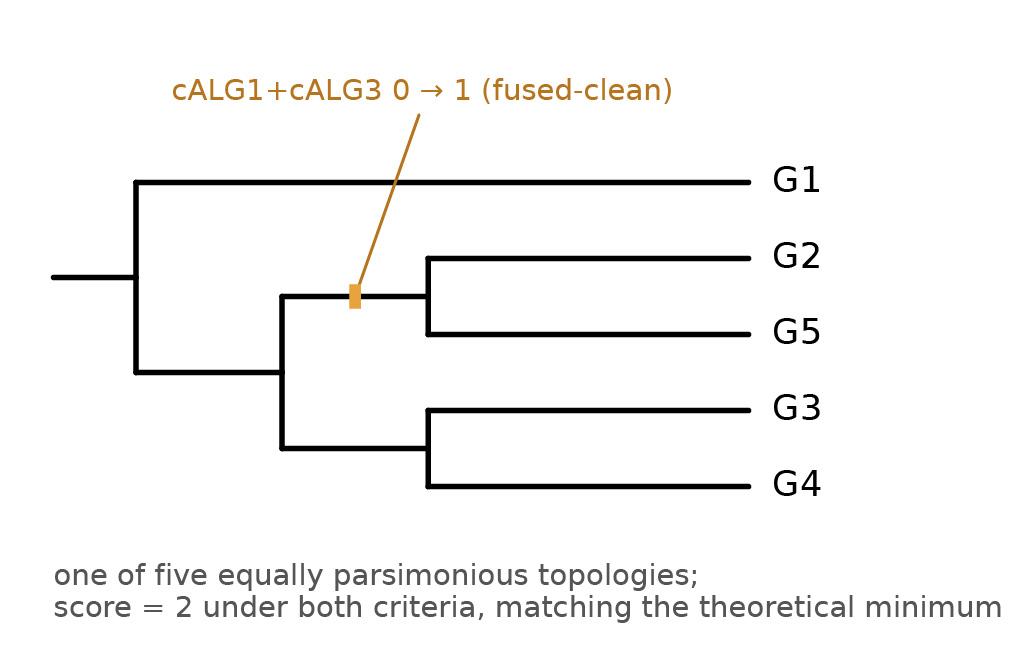

**Figure S3.** Rendered cladogram for the five-genome synthetic dataset under the reversible criterion, one of five equally parsimonious topologies. Both criteria returned a minimum score of 2, matching the theoretical lower bound of one fewer step than the number of observed states for a ternary character, which provides an independent check on the Sankoff scoring. With only one informative character the data cannot resolve five taxa, and no bipartition beyond the rooting is shared by all five topologies; Anvaya reports the tie count and details one representative topology. The synapomorphy uniting G2 and G5 in this topology, an independent gain of the clean fusion between the first two linkage groups, is labelled on the branch where it originates.

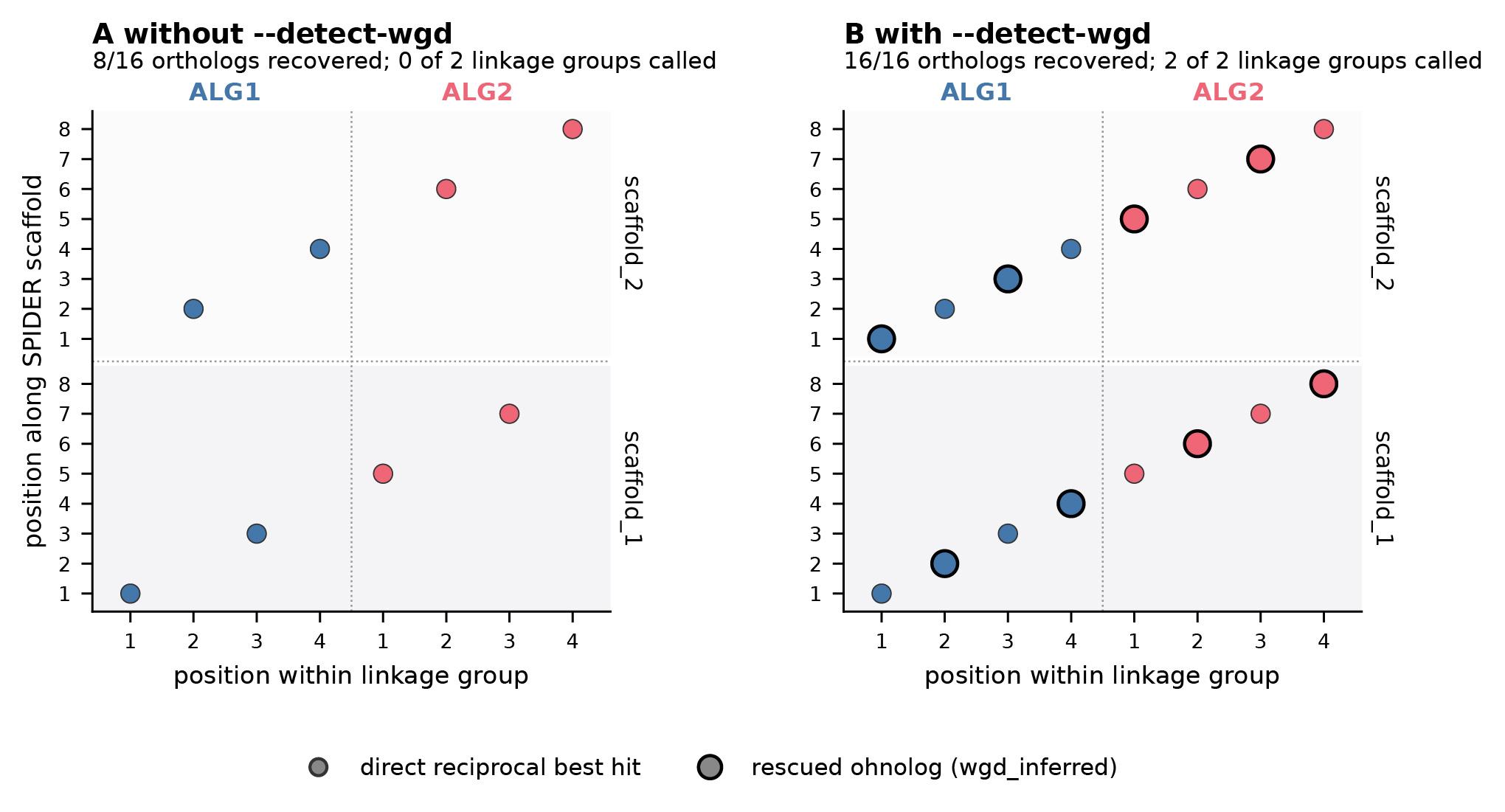

**Figure S4.** Oxford-style synteny dot plots of the simulated duplicated genome SPIDER against the non-duplicated genome TICK. (A) Without ohnolog rescue, plain reciprocal best hit recovers only 8 of 16 true orthologs, split thinly across both ohnologous scaffolds, and neither linkage group clears the significance threshold on either scaffold, so the duplication is invisible rather than conspicuous. (B) With --detect-wgd, all 16 orthologs are recovered and both linkage groups are correctly called, in full, on both ohnologous scaffolds. Compare Table S2.

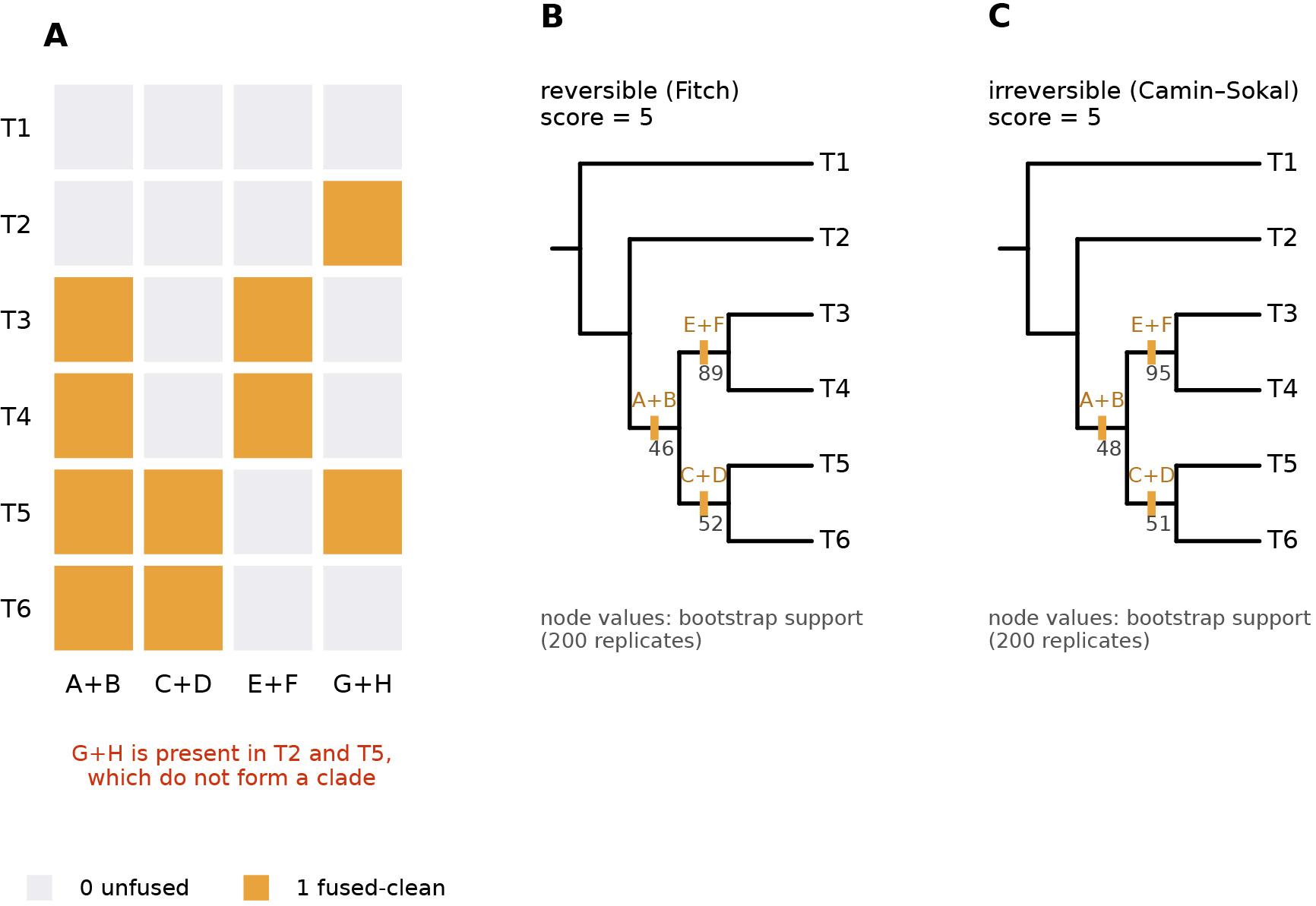

**Figure S5.** Six-taxon diagnostic dataset. (A) Character-matrix heatmap: the three clean, nested synapomorphies (A+B, C+D, E+F) form contiguous blocks corresponding to real clades, while the deliberately homoplastic G+H character is scattered across T2 and T5, which do not form a clade — the pattern the consistency index is designed to flag. (B) Most-parsimonious tree under the reversible criterion and (C) under the irreversible criterion. The exhaustive search recovered a single tree, (T1,(T2,((T5,T6),(T3,T4)))), the designed true tree, with a score of 5 under both criteria, and mapped all three designed synapomorphies onto exactly the clades built into them. Bootstrap support (200 replicates) was 89% (reversible) and 95% (irreversible) for the E+F-supported clade (T3,T4), against 46–52% for the two clades whose support competes with the homoplastic character. The permutation test gave p = 0.309 (reversible) and p = 0.060 (irreversible) over 200 replicates, with mean shuffled scores of 5.69 and 6.76 against an observed 5: the real matrix is more parsimonious than its shuffled counterparts, but at four characters the test cannot reach conventional significance, the correct result for a matrix this small.
